# Olfactory detection of viruses shapes brain immunity and behavior in zebrafish

**DOI:** 10.1101/2023.03.17.533129

**Authors:** Aurora Kraus, Benjamin Garcia, Jie Ma, Kristian J. Herrera, Hanna Zwaka, Roy Harpaz, Ryan Y. Wong, Florian Engert, Irene Salinas

## Abstract

Olfactory sensory neurons (OSNs) are constantly exposed to pathogens, including viruses. However, serious brain infection via the olfactory route rarely occurs. When OSNs detect a virus, they coordinate local antiviral immune responses to stop virus progression to the brain. Despite effective immune control in the olfactory periphery, pathogen-triggered neuronal signals reach the CNS via the olfactory bulb (OB). We hypothesized that neuronal detection of a virus by OSNs initiates neuroimmune responses in the OB that prevent pathogen invasion. Using zebrafish (*Danio rerio*) as a model, we demonstrate viral-specific neuronal activation of OSNs projecting into the OB, indicating that OSNs are electrically activated by viruses. Further, behavioral changes are seen in both adult and larval zebrafish after viral exposure. By profiling the transcription of single cells in the OB after OSNs are exposed to virus, we found that both microglia and neurons enter a protective state. Microglia and macrophage populations in the OB respond within minutes of nasal viral delivery followed decreased expression of neuronal differentiation factors and enrichment of genes in the neuropeptide signaling pathway in neuronal clusters. Pituitary adenylate-cyclase-activating polypeptide (*pacap*), a known antimicrobial, was especially enriched in a neuronal cluster. We confirm that PACAP is antiviral *in vitro* and that PACAP immunoreactivity increases in the OB 1 day post-viral treatment. Our work reveals how encounters with viruses in the olfactory periphery shape the vertebrate brain by inducing antimicrobial programs in neurons and by altering host behavior.

## Introduction

The nervous and immune systems work in tandem to recognize, escape, and eliminate pathogens (Kraus et al., 2021). Even in basal metazoans like ctenophores and cnidaria, bidirectional neuroimmune communication in specialized sensory cells influences motility and immunity (Babonis et al., 2018; Klimovich et al., 2020; Kraus et al., 2021; Pita et al., 2018; Rivera et al., 2012; Varoqueaux et al., 2018). In vertebrates, pathogens can influence immune responses by directly activating sensory neurons (Chiu et al., 2013; Pinho-Ribeiro et al., 2023; Sepahi et al., 2019). In mammals, sensory neurons at mucosal barriers are heavily studied because of their close proximity to pathogens, their ability to transmit pathogen signals from the periphery to the brain, and their potent immune regulatory functions. As such, sensory neurons found at mammalian mucosal barriers are the target of therapeutic interventions from pain and itch to infectious diseases (Chesne et al., 2019; Godinho-Silva et al., 2019; Huh and Veiga-Fernandes, 2020; Pinho-Ribeiro et al., 2017).

Chemosensory olfactory sensory neurons (OSNs) in the nasal mucosa directly contact the environment, relay information to the brain, and shape animal behavior from feeding to mating and predator avoidance (Curtis, 2014; Padilla-Coreano et al., 2016; Padilla-Coreano et al., 2019). Exposure to the environment leaves OSNs, and the olfactory bulb (OB) they project to, vulnerable to viral infection (Mori et al., 2002; Moseman et al., 2020; van Riel et al., 2015; Wheeler et al., 2017). Despite this exposed localization, viral infection of the CNS via the olfactory route is rare (Butowt et al., 2021). This is because OSNs and other cells in the nasal mucosa coordinate rapid and effective antiviral immune responses to staunch viral spread (Ai and Klein, 2020; Bi et al., 1995; Durrant et al., 2016; Kurhade et al., 2016; van den Pol et al., 2014). In support, disruption of the olfactory epithelium and destruction of OSNs results in impaired antiviral immunity in the OB, indicating that neurons play an important role in protection of the CNS (van den Pol et al., 2014).

Neuroimmune interactions along the olfactory-brain axis are often studied during active infection of neurons. In the OB, neurons can directly detect vesicular stomatitis virus (VSV) via pattern recognition receptors and secrete chemokines to recruit immune cells necessary for viral clearance (Ghita et al., 2021). Furthermore, microglia in the OB act as antigen presenting cells to T cells in the VSV infection model (Moseman et al., 2020). Overall, the OB has been proposed to be an immuno-sensory region of the CNS based on evidence from mouse models where viruses actively infect the OB (Durrant et al., 2016). However, how the OB responds to presence of viruses in the periphery is not well understood.

Neuropeptides are small chemical messengers released by neurons. Beyond their well-known neurotransmitter functions, neuropeptides link innate immunity and the nervous system via potent antimicrobial functions (Brogden et al., 2005). Antimicrobial activity in neuropeptides is an ancestral feature of the metazoan tree of life as illustrated by the diversity of *Hydra* neuropeptides which shape the microbiome via their antibacterial properties (Augustin et al., 2017; Klimovich et al., 2020). Other examples of neuropeptides with known antimicrobial activities include substance P, neuropeptide Y (NPY) or pituitary adelylate cyclase activating polypeptide (also known as PACAP) (Brogden et al., 2005). In both invertebrates and vertebrates, the olfactory system expresses several neuropeptides (Hansel et al., 2001; Hussain et al., 2016). Neuropeptide functions in olfactory circuits include neuromodulation, regulation of neurogenesis, learning and behavior (Chalasani et al., 2010; Hansel et al., 2001; Hussain et al., 2016; Lindsay et al., 2022; Negroni et al., 2012) Yet, how pathogens reaching the olfactory system impact neuropeptide expression and whether this is a mechanism for pathogen control is currently unknown.

Neurons can transmit the presence of a pathogen within milliseconds and rapidly release soluble mediators, thereby increasing velocity of immune responses (Cohen et al., 2019). We have reported that the presence of a live rhabdovirus, infectious hematopoietic necrosis virus (IHNV), in the olfactory organ of teleost fish can generate neuronal signals and very rapidly induce expression of select immune transcripts in brain tissues (Sepahi et al., 2019). We previously demonstrated that in rainbow trout, this occurred largely due to signals initiated by crypt neurons, a specialized type of neurons with S100 and TrkA immunoreactivity (Ahuja et al., 2013). Our studies also suggested that multiple OSN types beyond crypt neurons respond to olfactory viruses based on pERK stainings (Sepahi et al, 2019). However, how these ultra-fast immune responses impact neuronal states in the brain, specifically in the OB, and whether they are integrated into behavioral responses is currently unknown. Here we present a zebrafish model to investigate the neuroimmune interactions that occur in the OB of zebrafish in response to presence of a virus in the periphery.

## Results

In rainbow trout, intranasal (I.N.) delivery of IHNV causes apoptosis in a subset of OSNs in the olfactory organ (OO) as rapidly as 15 minutes (m) as the lymphocyte cellular landscape and chemokine expression profile changes in the OB (Sepahi et al., 2019). While live attenuated IHNV is present in the olfactory organ of rainbow trout until 4 days (d) post I.N. delivery, viral RNA is never detected in the brain (Larragoite et al., 2016). Here we adapted this model to adult zebrafish. Virulent IHNV has been used to infect zebrafish before by adapting the virus to the warmer host temperatures (Aggad et al., 2010; Levraud et al., 2014; Ludwig et al., 2011). However, we aimed to restrict the presence of the virus to the olfactory organ and avoid penetration of the brain. Using I.N. delivery of live attenuated IHNV at the normal host temperature (28), we measured the kinetics of the viral detection in the OO and OB (Figure 1). By qPCR we only detected RNA of the IHNV nucleoprotein 15 m after I.N. delivery, and it was never detected in the OB (Figure 1B). This confirmed that in our model IHNV is only transiently present in the olfactory periphery and does not reach the OB.

**Figure 1.**
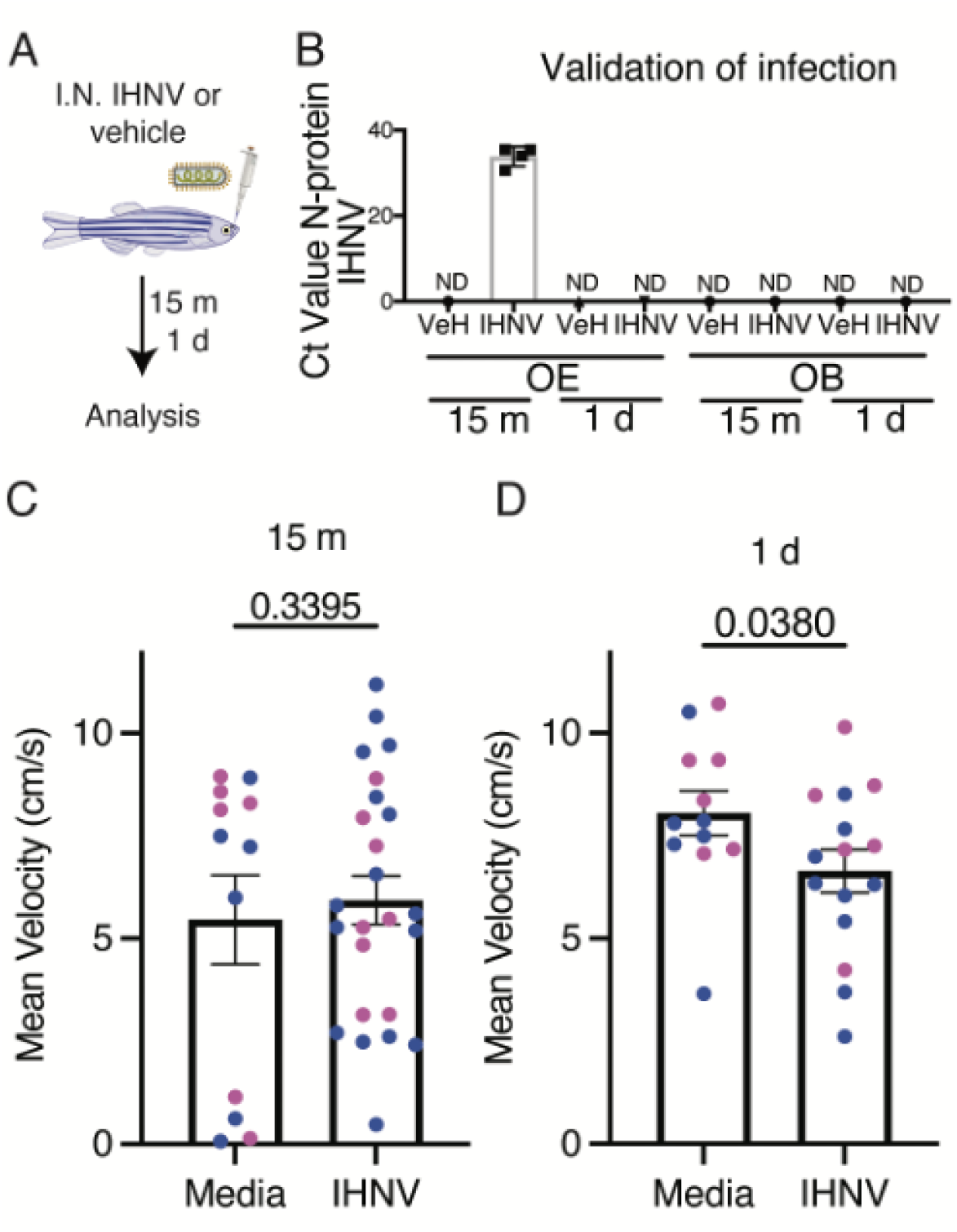
Without viral translocation to the brain I.N. IHNV changes adult behavior at 1. **d.** (A) schematic of intranasal administration of aquatic rhabdovirus infectious hematopoietic necrosis virus (IHNV) or vehicle (VeH). (B) quantification of IHNV N-protein presence in the olfactory epithelium (OE) and olfactory bulb (OB) 15 m and 1 d after I.N. delivery of virus. N = 4-6, each replicate is a pool of 4 animals. (C) Average velocity of the animal over the test for 15 m (C) or 1 d (D) after IHNV or media control. Test was run for 5 m after IHNV or control odorant was added to the flow cell. Statistics are Student’s T test. Error bars are SEM. Male (blue) and female (pink) animals do not show a difference in behavior.

### Detection of live attenuated IHNV triggers behavioral changes in adult and larval zebrafish

We wanted to determine if zebrafish respond behaviorally to live virus. First, we tested adult animals I.N. treated with IHNV or vehicle controls and monitored swimming behavior in an open field paradigm 15 m or 1 d post-treatment (Baker et al., 2018). We observed significant behavioral changes in adult animals at 1 d, but not 15 m. Specifically, treated zebrafish had decreased mean velocity, potentially indicative of classical sickness behavior (Figure 1C-D).

We also investigated if 6 day post-fertilization (dpf) larvae also exhibit behavioral changes to live viral detection. In this model, larvae were exposed to the virus or a vehicle in the water for 30 seconds and swimming behavior was tracked for 22 m. When exposed to IHNV, larva swam at significantly lower velocities, and displayed a trend towards increased turning behavior (Figure 2A-B) compared to the vehicle. Decreased velocity in larvae is a known stress response (Lopez-Luna et al., 2017). Together, our results indicate that olfactory detection of IHNV by both adult and larval zebrafish is integrated in higher areas of the brain and translates into behavioral responses.

**Figure 2.**
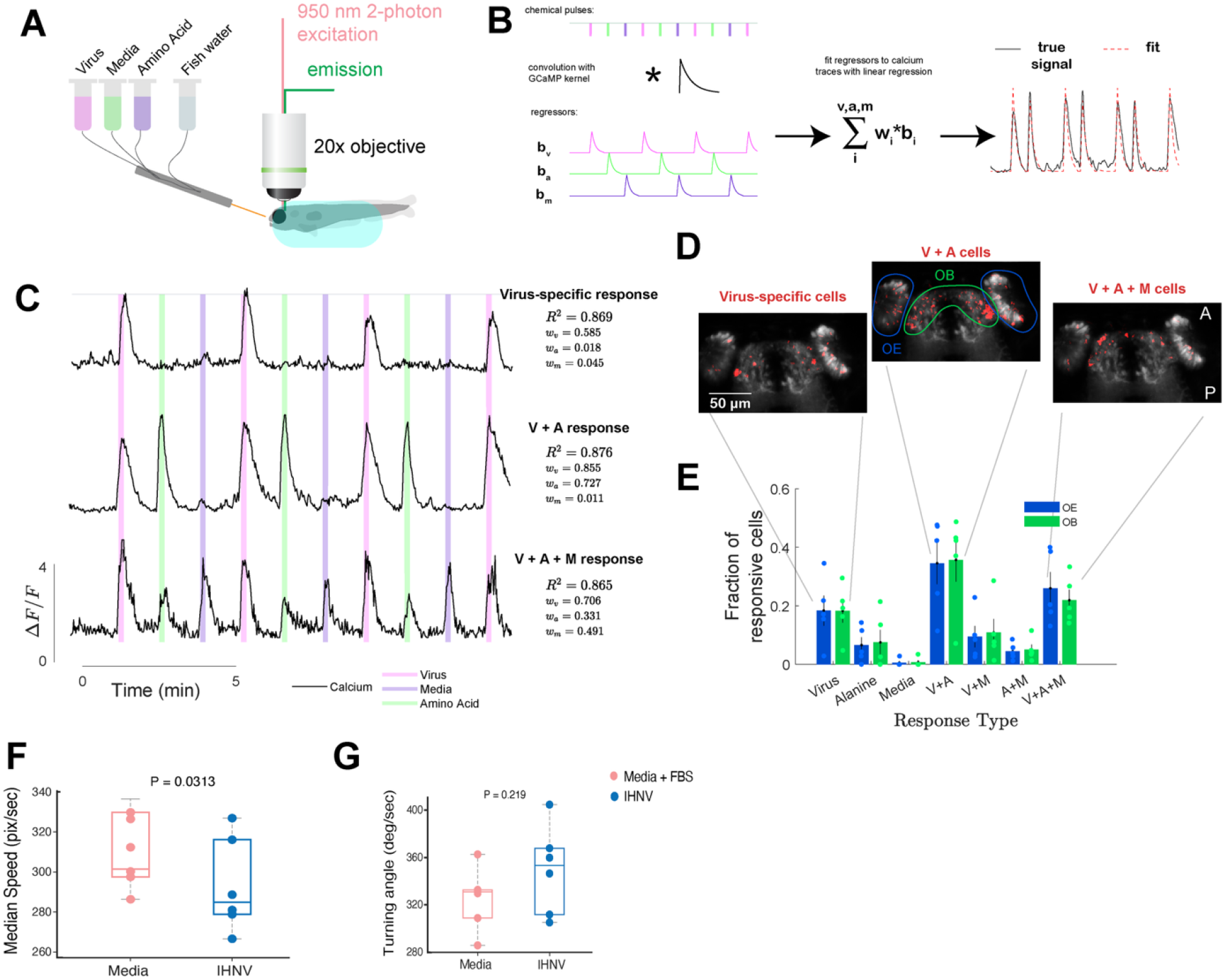
Neuronal activation of the zebrafish OO and OB and behavioral changes in response to nasal viral delivery. Schematic of 2-P imaging preparation with larval zebrafish embedded in 2% agarose and nose exposed for delivery of different chemical cues (A). Schematic of regression approach to identify response types (B). Briefly, the timings of the stimulus pulses - virus (pink), amino acid (green), and control media (blue) are convolved with a kernel representing the GCaMP6S dynamics to generate regressors for each stimulus. Linear regression is then used to fit linear combinations of these regressors to calcium signals which yield weights *w* that can be used to classify cell types as in (C). Example calcium traces of olfactory epithelial neurons that illustrate the three most common response types (Virus-specific, virus (V) and amino acid (A) responsive, and all-tested chemical responsive (A+M+V)) (C). The generated fit and regressor weights are provided to the right of each trace. Example locations within the olfactory epithelia and OB of the three most common cell types (D). Each image is a maximum projection of the anatomy and cell location. Cells that that belong to the given class are labelled in red. Bar graph depicting the average fraction of cells across fish (n=5) with a good fit (90^th^ percentile R^2^ within each fish) that belong to each possible response type (E). Blue indicates fraction of responsive cells within the OB, green bars indicate fraction of cells within the bulb. (F) Median speed of larva in a lateral flow chamber exposed to IHNV for 4 seconds. Speed adjusted for the flow rate. (G) Turning angle of larval zebrafish in lateral flow cell. Test was run for 5 m after IHNV or control odorant was added to the flow cell. Statistics for behavioral assays are Student’s T test. Dots represent individual fish. OE, olfactory epithelia; OB, olfactory bulb.

### Viral-specific activation of neurons in the OO and OB of larval zebrafish

In the rainbow trout live attenuated IHNV causes action potentials in the OSNs, as measured by EOG (Sepahi et al., 2019), but electrical responses in the teleost brain were not recorded. Because I.N. IHNV caused behavioral changes in adult and larval zebrafish, we asked whether live attenuated IHNV could activate OSNs and consequently neurons within the OB.

We live imaged Tg(vGlut2a:Gal4)^nns20^;Tg(UAS:GCaMP6S)^13a^ 5 dpf larva by 2-P microscopy and recorded calcium-dependent fluorescent signals in response to attenuated IHNV, control media, or an amino acid, alanine (Figure 2A).

Using stimulus-specific regressors (Figure 2B, methods) we identified cells that were specific to individual cues or combinations. Results show the presence of regions of interest (ROIs) in both OSNs and the OB which are specifically activated by IHNV but not alanine (positive control) or culture medium + 5% FBS (vehicle control for IHNV) (Figure 2C-E & Figure S1). As expected, there were also ROIs only activated by alanine, or combinations of the three odorants which a widespread localization in the epithelia and OB compared to IHNV-specific cells (Figure 2E & Figure S1). These results reveal populations in the OB activated by transient interactions with IHNV in the epithelia concordant with behavioral changes.

### scRNA-Seq of the zebrafish OB after intranasal IHNV reveals changes to the cellular landscape

Given observed neuronal activation of the OB and behavioral changes, we sought to identify potential molecular changes in the OB at the single cell level following I.N. IHNV delivery. While scRNA-Seq has been performed on the OB in a murine model of olfactory occlusion (Tepe et al., 2018), comprehensive single cell transcriptomic studies have not been performed in the adult zebrafish OB. We identified a total of 4 lymphocyte clusters, 8 microglia clusters, 5 neuronal clusters, and 2 oligodendrocyte clusters integrating vehicle and IHNV treated datasets (Figure 3A-B).

**Figure 3.**
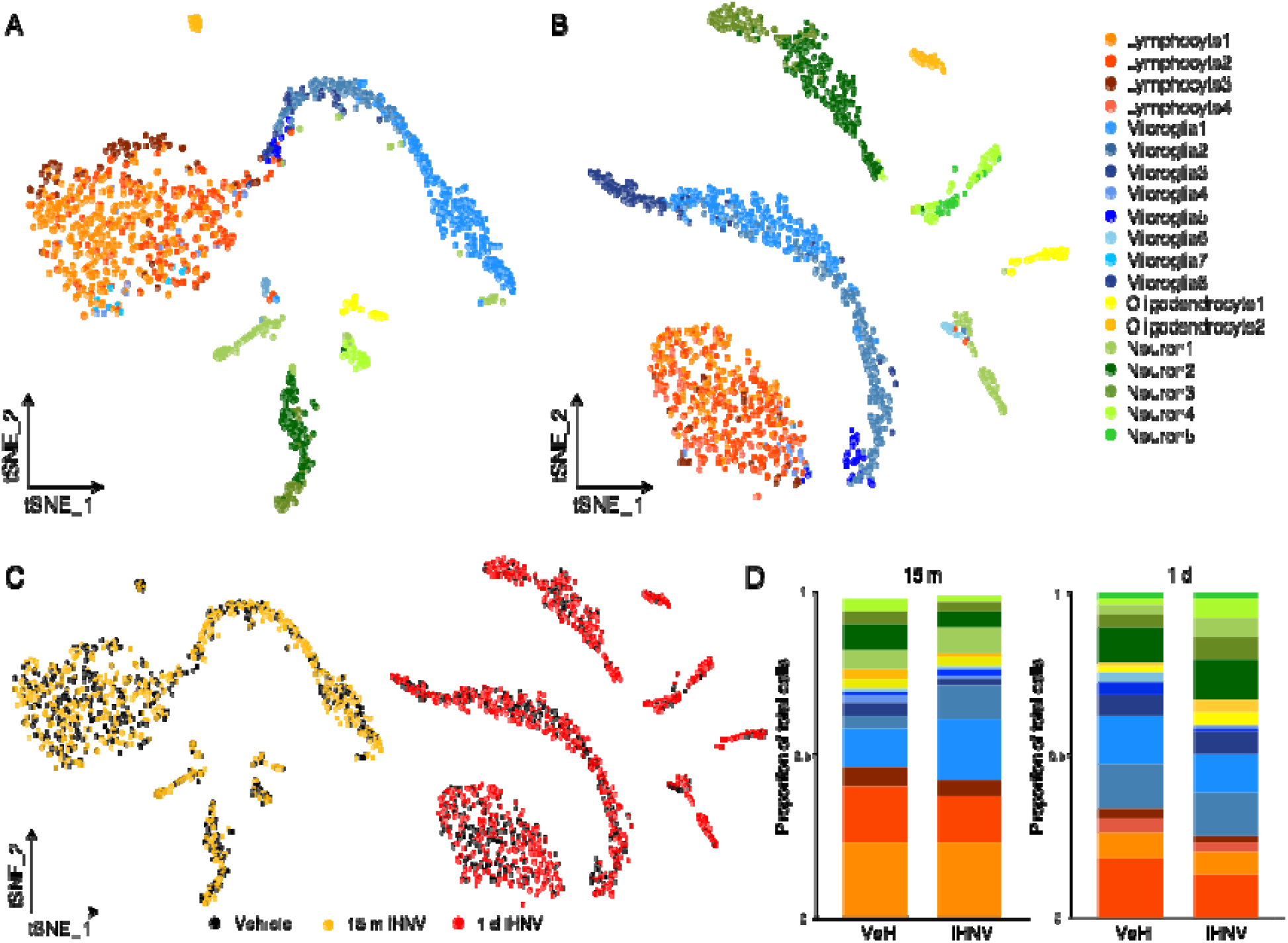
sc-RNA seq of the OB reveals changes to the cellular landscape after I.N. IHNV. tSNE plots of single cells isolated from the adult zebrafish OB at 15 m (A) and 1 d (B). (C) tSNE plots showing Vehicle (VeH) (black), 15 m IHNV (yellow), and 1 d IHNV (red). (D) Bar plots of proportions of cells from each cluster for each treatment. Statistical proportional analysis for changes to the proportions following IHNV treatment is in supplemental figure 3.

Microglia1, Microglia2 and Microglia4 constitute the majority of microglia sequenced (Figure S2) and express classical microglial markers (*apoeb, mpeg1.1,* and *marco*) in addition to markers for mobile, ameboid and phagocytic microglial phenotype (*ccl34b.1* and *cd63*) (Figure S2&S4) (Mathys et al., 2017; Silva et al., 2021; Wu et al., 2020). At 15 m post I.N. IHNV treatment, Microglia1 and Microglia2 proportions both significantly expand relative to the 15 m I.N. controls (Figure 3C-D & S3A-C) suggesting a quick increase in activated microglia following I.N. IHNV delivery. Gene ontology (GO) of differentially regulated genes from these rapidly expanding microglial clusters at 15 m after virus compared to vehicle treated microglia shows enrichment of positive regulation of nervous system development and response to hypoxia. Hypoxia response was particularly interesting as it has been related to antiviral defense (Huang et al., 2021) (Figure S4).

Microglia3 express many of the same markers as Microglia1-2 but have higher expression levels of MHC II features such as *cd74a* and *cd74b* suggesting they are professional antigen presenting cells (APCs) (Figure S2&S4). Microglia3 and 5 also express a separate subset of known zebrafish microglial markers but may represent infiltrating macrophage phenotypes since they express *ccl19a.1* and *siglec15l* (Figure S2&S4) (Oosterhof et al., 2018). These clusters also express anti-inflammatory *tnf-b* and *grn2* (Figure S4). Further, Microglia4 and Microglia8 express *il4* and *il13,* characteristic of a type II, anti-inflammatory state (Quarta et al., 2020). Conversely, Microglia6 express pro-inflammatory markers (*mmp2* and *mmp14*) (Figure S2) (Konnecke and Bechmann, 2013). At 1 d after I.N. IHNV treatment, changes in microglia clusters were less profound than at 15 m. At 1 d, *mmp2* expressing Microglia6 is expanded (Figure 3C-D & S3A-C). This is particularly interesting as *mmp2* has been shown to affect function of neuronal progenitors *in vitro* (Li et al., 2017). Lastly, Microglia8 is a small cluster characterized by expression of thus far unannotated genes (Figure S2). Combined, these data show that OB microglia mount rapid and complex immune responses to peripheral viruses in zebrafish.

We previously reported that CD8^+^ T cells migrate into the OO of rainbow trout 15 m after I.N. IHNV concurrent with an egress of CD8^+^ T cells from the OB (Sepahi et al., 2019). Moreover, we found that the cellular landscape of lymphocytes changes after I.N. SARS-CoV-2 S RBD in the zebrafish OO (Kraus et al., 2022). These findings made us look for lymphocyte clusters and their responses in the zebrafish OB scRNA-Seq dataset (Figure S5). One caveat to this data set is that animals were not perfused and therefore lymphocytes residing in the OB vasculature were likely included in the OB single cell suspensions. Further, the medium used during extraction was optimized for neuronal survival rather than viability of immune cells and consequently the lymphocyte clusters did not have high sequencing depth. Despite these caveats we captured 4 lymphocyte clusters.

In the vehicle treated OB, approximately 20% of collected cells clustered as Lymphocyte1 and 15% of cells clustered as Lymphocyte2 (Figure S5A). Further, approximately 5% of cells were B cells clustered in Lymphocyte3 (Figure S5A). Lymphocyte1 cells expressed many markers associated with NK cells in zebrafish (*ccl38.6, ccl34b.4,* and *ccl38a.5*) as well as multiple novel immune type receptors (*nitr*) thought to function as natural killer (NK) cell receptors and are therefore an innate-like cluster (Figure S2) (Carmona et al., 2017; Wei et al., 2007). Lymphocyte2 cells express classical markers of T cells including *cd4*, *cd8,* and *cxcr4* (Figure S2). Interestingly, 1 d after I.N. IHNV, a fourth cluster of lymphocytes emerged (Figure S3). Lymphocyte4 expresses *cd4* as well as *foxp3a,* the master transcription factor and marker for T_regs_ (Morikawa and Sakaguchi, 2014) (Figure S2). While the lymphocyte populations do not change within timepoints, at 1 d Lymphocyte2 (classical T cell cluster) becomes the dominant cluster rather than innate-like cluster Lymphocyte1 which is the larger lymphocyte cluster in the integrated 15 m samples (Figure S3).

Re-clustering cells from Lymphocyte1, Lymphocyte2, and Lymphocyte4 with 5 significant principal components (PCs), reveals 6 distinct populations of non-B cell lymphocyte clusters (Figure S5). While all clusters express potential zebrafish NK markers (*ccl38.6* and *ccl34b.4*) and to a lesser extent *nitrs,* there are some differences across clusters (Figure 3C). For example, only two clusters express apoptosis-inducing serine kinase *gzm3.4,* indicating they are cytotoxic. Interestingly, neither of these clusters express *cd8a* or *cd8b*, and one of them expresses *rorc,* the gene for transcription factor Ror-gt, indicating that these clusters are likely cytotoxic NK cells (red) and ILC3 (mint green) (Figure S5C). One cluster (purple) expressed *il4*/*il13* and therefore may be ILC2s (Figure S5C). Importantly, there is one cluster (grass green) that expresses both *cd4* and *foxp3a* indicating that they are T_reg_ cells. Overall, these ILC-like expression profiles like those of zebrafish gut ILCs (Hernandez et al., 2018).

Within the neuronal clusters, neuron2 and neuron3 appear to be mature neurons expressing markers such as *pcdh7a, pcdh17,* and *olfm1b* (Figure S2) (Kanageswaran et al., 2015). Neuron3 are likely granule cells based on their expression of glutamate receptors (*grm1a* and *grm2a*) and GABA transport components (*slc32a1*) (Nagayama et al., 2014) (Figure S2). Overall, previous NGS endeavors have uncovered few genetic markers for mitral cells in mice, and almost none of these were found within our zebrafish mature neuron clusters (Tepe et al., 2018; Zeppilli et al., 2021). Classical mitral cell markers *cdhr1a* and *tbx21* were not detected in our data set, but we hypothesize that Neuron2 represent mature mitral cells based on glutamate associated genes present (*gad* and *gad1b*) (Bohm et al., 2020; Nagayama et al., 2014; Zeppilli et al., 2021) (Figure S2).

Neuron1 is an interesting cluster expressing immune related features (*ifn-g* and *ptgdsb.1*) in addition to an aquaporin gene (*aqp1a.1*) that has been linked to adult neurogenesis (Figure S2) (Zheng et al., 2010). Interestingly, IFNg is expressed in axonimized motor neurons and is important in differentiation and neurogenesis in the murine adult subventricular zone (Olsson et al., 1989; Pereira et al., 2015). Further, Neuron1 cells express many factors associated with neuronal neurogenesis (*nog2*, *mdka*, *notch3, sox2* and *her6*) and are likely progenitors recently migrated from the subventricular zone (Figure S2) (Lubke et al., 2022; Marz et al., 2010; Peretto et al., 2004; Rieskamp et al., 2018; Schmidt et al., 2013). While Neuron4 and Neuron5 clusters express markers of progenitor neurons, they also have markers that suggest they are further along in differentiation programs such as *lhx9*, *neurod1* and *gap43* (Figure S2) (Schmidt et al., 2013). Neuron5 was only found in the 1 d IHNV data set, indicating an increase in the proportion of differentiating neurons that have gained a unique phenotype from viral exposure (Figure S3). After 1 d I.N. IHNV Neuron1 and Neuron5 have increased in proportion relative to the control cell suspensions (Figure 3C-D & S3A-C). There were 2 clusters of oligodendrocytes each expressing markers (*olig2, olig1,* and *sox10*) (Figure S2) (Takada et al., 2010; Tepe et al., 2018).

### Transcriptional profiles of OB neurons reveal increased developmental program at 1 d after intranasal delivery of virus

1 d after I.N. IHNV we observed many differentially regulated features pertaining to neurogenesis and differentiation. In mature Neuron3 *ucp2* is significantly upregulated, and cells in progenitor like clusters Neuron1 and Neuron5 also express more *ucp2* (Figure 4A-C). Murine studies showed *ucp2* to be necessary for normal proliferation and differentiation in embryonic neurogenesis (Ji et al., 2017) and UCP2 is an important regulator of innate immune responses via mROS inhibition (West et al., 2011). Neuron2 has upregulated *nrn1a,* known to facilitate neurite growth, and *irs2b,* important for neuronal proliferation during brain development and involved intracellular insulin signaling (Schubert et al., 2003; Shimada, 2013; Shimada et al., 2016) (Figure 4A-C). Neuron5 significantly downregulates *tac1,* which can be alternatively spliced into substance P and neurokinin A and is known to be expressed in the zebrafish olfactory system and inhibit olfactory stimulus in glomeruli across the animal kingdom suggesting a release of repressive signals (Figure 4A-C, E) (Nassel et al., 2019; Ogawa et al., 2012). Further, Neuron5 has significantly up-regulated *her15.2* and *pax6* which have both been associated with axonal growth in adult neurons (Figure 4A-B) (Mizoguchi et al., 2020; Sebastian-Serrano et al., 2012). Neuron1 and Neuron4 clusters have upregulated many transcription factors involved in differentiation (*nfia, zic2, sox6,* and *zfhx4*) and plasticity (*nfil3*) of neurons (Batista-Brito et al., 2009; Hemmi et al., 2006; Plachez et al., 2012; Tiveron et al., 2017; van der Kallen et al., 2015) (Figure 4A-C). Gene ontology of differentially expressed genes across neuronal clusters 1 d after I.N. IHNV relative to controls show biological processes such as neuropeptide signaling, CNS and glial development, and cell differentiation are enriched after viral treatment (Figure 4D) (Zhou et al., 2019). Overall, these data indicate that zebrafish OB neuronal clusters are increasing expression of genes with predicted functions in neurogenesis 1 d after I.N. IHNV treatment relative to control treated neurons.

**Figure 4.**
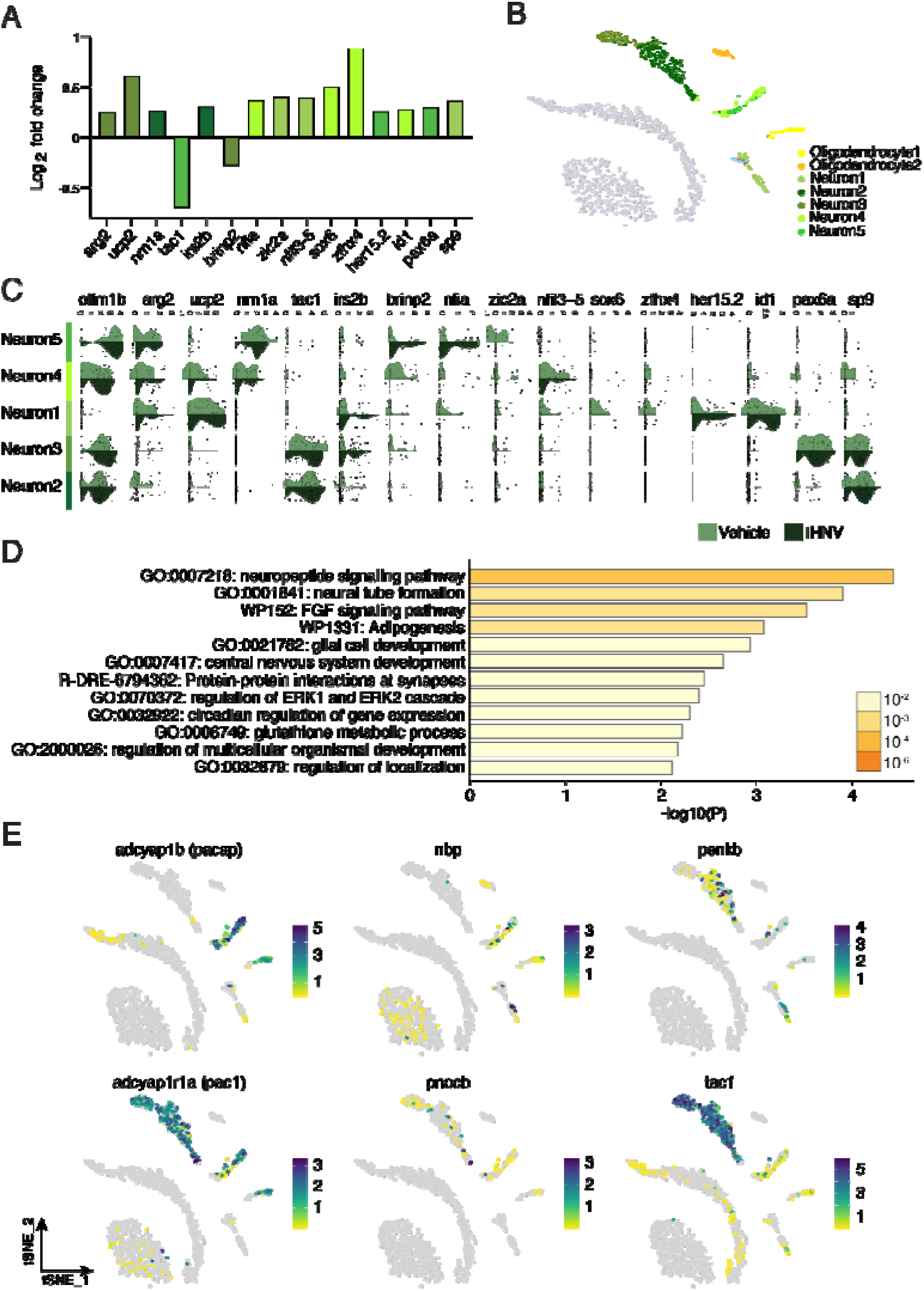
Transcriptional profiles of neurons reveal increased development at 1 d afte intranasal delivery of virus in adult zebrafish. (A) Bar plots showing fold change expression of significantly differentially expressed genes in neurons 1 d after I.N. IHNV relative to control. Bars are colored according to the cluster in which they are differentially expressed. (B) Tsne highlighting only the neuronal clusters (C) violin plots of differentially expressed genes in neuronal clusters 1 d after intranasal IHNV delivery shows many genes related to plasticity and development of neurons are upregulated compared to vehicle treated animals. (C) gene ontology of all differentially expressed genes in neuronal clusters 1 d I.N. IHNV compared to 1 d I.N. vehicle treatment. Gene ontology was run on Metascape (Zhou et al., 2019).(E) Feature plots for genes enriched in the top GO:0007218 Neuropeptide signaling pathway (*pacap*, *npb*, *penkb*, *scg5*, and *tac1*) and pacap receptor *pac1*.

To understand our scRNA-Seq data we performed *in vivo* Edu labeling of proliferating cells which may explain the increases in immature neuronal populations in response to virus. No increase in proliferation in the OB 1 d after I.N. IHNV was found (Figure S6). Therefore, while there is an increase in progenitor-like neurons following I.N. IHNV treatment detected by scRNA-Seq, it is not due to proliferation within the OB.

The most highly enriched gene ontology pathway in neurons was the neuropeptide signaling pathway and highlighted genes such as *adcyap1b* (also known as *pacap*), *npb*, *penkb*, *scg5*, and *tac1* (Supplemental Table 1). Feature plots reveal that *pacap (adcyap1)*, an evolutionarily conserved neuropeptide (Cardoso et al., 2020), is specifically expressed in clusters Neuron4 and Neuron5, which are expanded 1 d after I.N. IHNV (Figure 4E & S3). *pacap* was expressed in microglia8 at low levels and was not found in any other immune clusters. Further, the PACAP specific receptor *adcyap1r11* (or *pac1*) was expressed in mature neuron clusters, while VIP receptors (*vipr1a*, *vipr1b*, and *vipr2*), which PACAP is also a ligand for, were not detected in our transcriptomes (Figure 4E). Because PACAP-38 has been shown to be antimicrobial in both teleost and mammals (Lee et al., 2021; Semple et al., 2019; Velazquez et al., 2020) and to regulate feeding behavior in fish (Lugo et al., 2008) we decided to investigate this neuropeptide further.

### PACAP immunoreactivity increases in the zebrafish OB 1 d after I.N. but not I.P. IHNV delivery

To confirm our scRNA-Seq results, we performed immunofluorescence staining with an antibody against PACAP-38, using a commercially available anti-mouse PACAP-38 antibody. We performed amino acid alignments of zebrafish and mouse PACAP and determined 89% identity between both sequences as previously described in other comparative studies (Figure S7). We compared zebrafish treated I.N. IHNV or vehicle 1 d following delivery because at this time point, we observed changes in both behavior and transcriptional profiles of neurons. We found that PACAP-38 immunoreactivity is significantly higher in the OB of I.N IHNV treated zebrafish compared to the OB of zebrafish which received vehicle (Figure 5A-C). The isotype control, using normal rabbit IgG, showed no immunoreactivity, and blocking of the anti-PACAP antibody with the fish PACAP-38 peptide inhibited immunofluorescent signal (Figure S8&S9). Since PACAP responses have been detected in the mouse brain in the context of systemic inflammation or infection (Lee et al., 2021), we compared PACAP responses in the OB of zebrafish I.P. injected with IHNV or vehicle at the same time point. In contrast with the I.N. treatment, I.P. injected animals had decreased PACAP-38 immunoreactivity in the OB compared to vehicle controls (Figure 5D-F). To determine whether increases in PACAP-38 immunoreactivity were due only to changes in inflammatory state, we performed the same experiment in *Tg(NFkB:EGFP)^nc1^*transgenic fish and stained for EGFP. The results for this staining showed no difference in EGFP immunoreactivity in the OB following I.N. vaccination, but did show an increase in this signal following I.P delivery, indicating there was no correlation, either positive or negative, between PACAP-38 immunoreactivity and OB inflammation (Figure S10). These results confirm that intranasal IHNV delivery triggers a neuronal PACAP response in the zebrafish OB.

**Figure 5:**
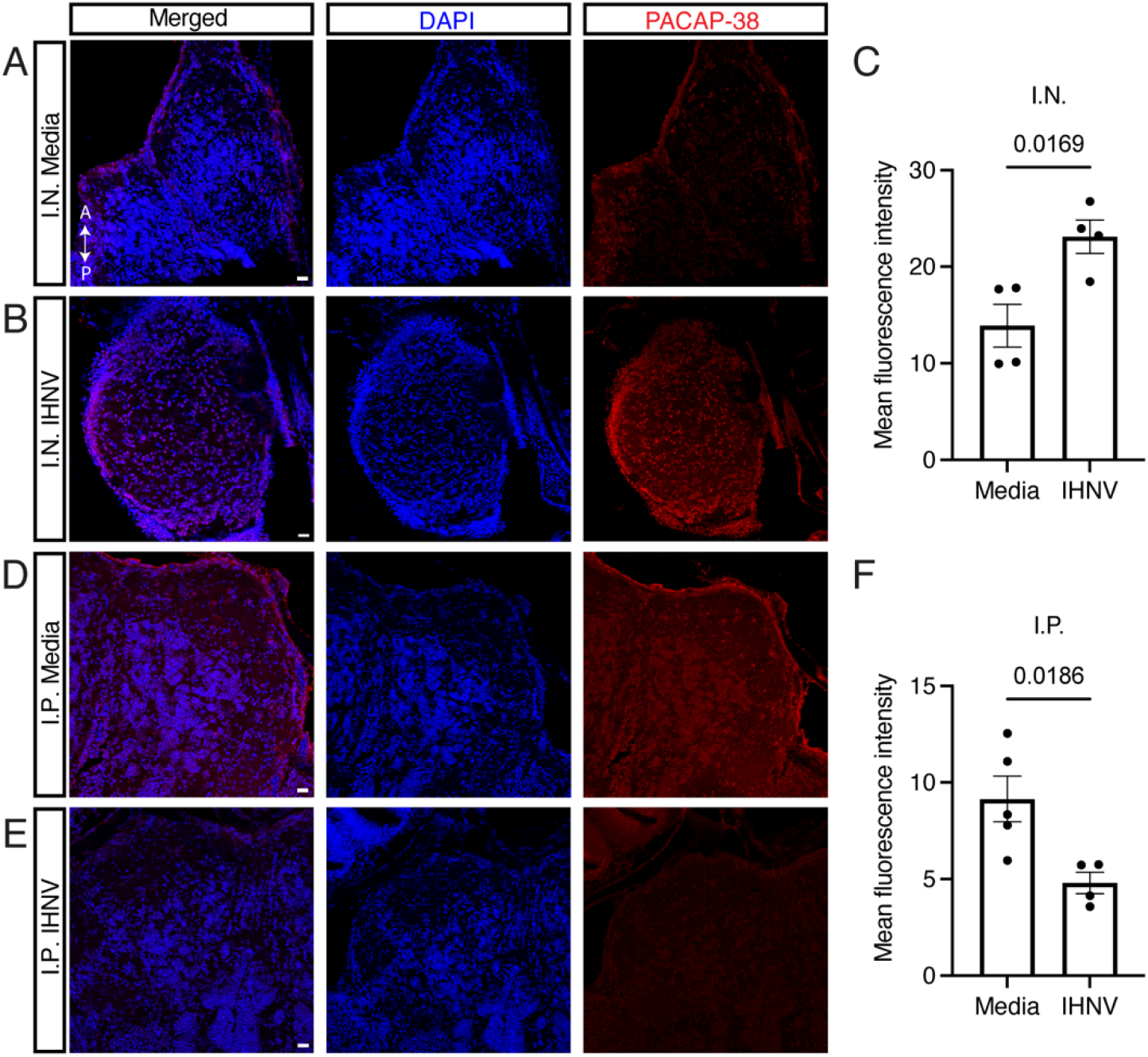
Neuropeptide PACAP-38 expression increases in the adult zebrafish OB 1 d after I.N. IHNV but not 1 d after I.P. IHNV treatment. (A) Representative confocal image of the zebrafish OB stained with anti-PACAP-38 antibody after receiving vehicle by I.N. delivery. (B) Representative confocal image of the zebrafish OB stained with anti-PACAP-38 antibody after receiving IHNV by I.N. delivery. (C) Quantification of the mean fluorescent intensity values for A and B. (D) Representative confocal image of the zebrafish OB stained with anti-PACAP-38 antibody after receiving vehicle by I.P. delivery. (E) Representative confocal image of the zebrafish OB stained with anti-PACAP-38 antibody after receiving IHNV by I.P. delivery. (F) Quantification of the mean fluorescent intensity values for D and E. P-values were calculated by a student’s T test, with error bars indicating standard error of the mean.

### PACAP-38 has antiviral activity against IHNV in vitro

To determine if the onset of *pacap* expression by immature neurons in the OB represents an antiviral immune program in zebrafish, we tested PACAP antiviral activity *in vitro*. We used recombinant amidated and non-amidated forms of PACAP-38 from the teleost *Clarias gariepinus* which has a 92% amino acid identity with *D. rerio* PACAP (Figure S8) (Lugo et al., 2008). Of note, amidated PACAP-38 is the natural, biologically functional form of PACAP found in vertebrates (Hansel et al., 2001). As an experimental control for recombinant protein cytotoxicity, we evaluated viability of teleost cell line Epithelioma papulosum cyprini (EPC), the standard method for propagating IHNV. Results from cell viability assays (Figure S11) show that the active amidated PACAP-38 was non-toxic at concentrations up to 100nM. At the lowest concentration of recombinant protein tested (10nM) for cytotoxicity, non-amidated PACAP-38 exhibited cellular toxicity with 85.6% EPC cell viability (Figure S11).

Next, we tested whether PACAP-38 (aminated and non-aminated) protects EPCs from cell death when infected with virulent IHNV. No antiviral effects were observed for the non-active form PACAP-38 at any concentration tested (Figure 6). The active form of amidated PACAP-38 provided protection from virulent IHNV infection in a dose-dependent manner (Figure 6). Up to 70% protection was achieved at the concentration of 1µM when the active PACAP-38 was added to the cells together with the virus during the infection period (competition assay) (Figure 6). EPC cells without PACAP-38 (either active or inactive) with IHNV showed typical morphologies of viral cytopathic effect (CPE). CPE was reduced at increasing concentrations of the active amidated PACAP-38 (Figure 6). Similar results were obtained in post-infection assays, where PACAP was added after the infection with IHNV (Figure 6).

**Figure 6.**
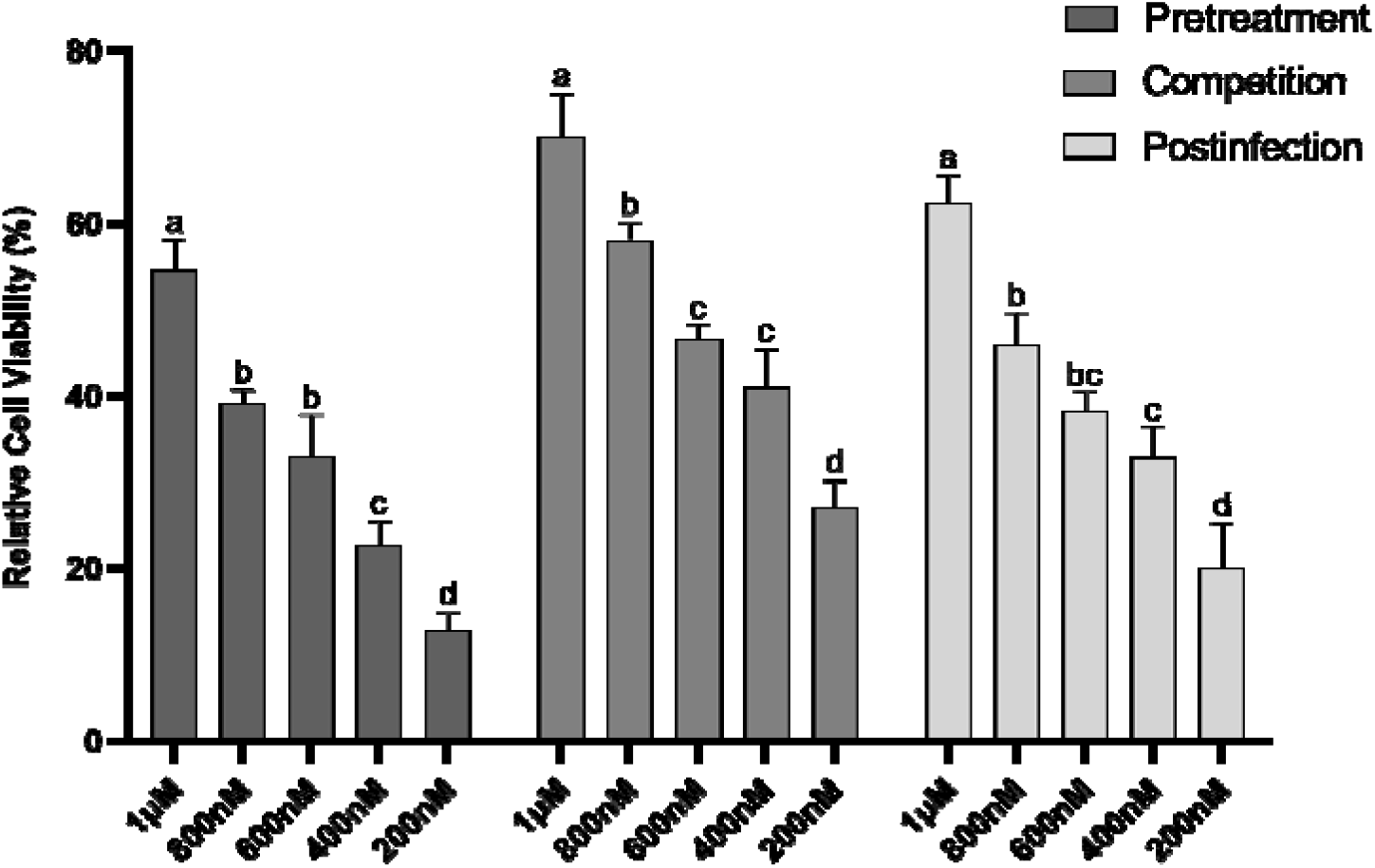
Amidated PACAP-96% protects EPC cells from IHNV-induced cell death. Relative percent viability of EPC cells that were either pretreated with amidated PACAP-38 prior to infection (pretreatment), treated with amidated PACAP-38 at the time of the infection (competition) or infected with IHNV and then treated with different concentrations of amidated PACAP-38 (postinfection). Results are representative of two independent experiments with N=4 for each treatment group. Different letters indicate significant differences among treatments by one-way ANOVA (P<0.05).

## Discussion

Chemosensory systems detect and respond to environmental dangers; the integration of these signals by the brain is essential for animal survival. The chemosensory olfactory system detects odorants and sends signals to higher regions of the brain via the OB. OSNs directly sense the environment and project their axons into the OB, making them ideal danger sensors (van Riel et al., 2015). Protecting the CNS from pathogen translocation falls to the OB, a dynamic region of the brain known to participate in immune defense in both mammals and fish (Bi et al., 1995; Christian et al., 1996; Durrant et al., 2016; Kalinke et al., 2011; Kurhade et al., 2016; Leyva-Grado et al., 2009). Immune responses in the OB have been described using murine models of neurotropic pathogen infection where pathogens could be detected in the OB following different routes of infection (Dinn, 1980; Flexner, 1936; Song et al., 2021; Torres-Fernández et al., 2018; van Riel et al., 2015; Wheeler et al., 2017). For example, I.N. infection with vesicular stomatitis virus leads to short infection of the OB without spread to the CNS. This is achieved by expression of type I interferons across the brain, which is abrogated when the olfactory epithelium is chemically ablated (van den Pol et al., 2014). However, immune responses in the OB caused by pathogen detection or damage in the olfactory periphery without translocation into the brain are not well understood.

Microorganisms and their products can directly interact and be recognized by neurons (Baral et al., 2016; Chiu et al., 2013; Pinho-Ribeiro et al., 2023). In teleosts, IHNV directly interacts with TrkA expressed on a subset of OSNs and triggers electrical signals in the olfactory epithelium (Sepahi et al., 2019). Furthermore, the teleost OB responds to the presence of virus in the peripheral OO without viral translocation by regulating innate immune gene transcription (Sepahi et al., 2019). It is unknown how rapidly neuroimmune responses in the OB prepare the brain for pathogen invasion. Here we present evidence in zebrafish that indicates that peripheral olfactory viruses trigger neuroimmune responses in the OB which are computed into behavioral change.

While some strains of IHNV can be neurotropic, in rainbow trout live-attenuated IHNV can only be detected in the OO up to 4 d after I.N. delivery and never enters the brain (LaPatra et al., 1995; Larragoite et al., 2016). In the present zebrafish model, live attenuated IHNV is only detectable in the OO of adult zebrafish 15 m after I.N. delivery and does not translocate to the OB. However, transient presence of a virus is enough to cause changes in behavior. Zebrafish are a powerful animal model in neuroscience in part due to behavioral features that can be used as a readout for human actions (Stewart et al., 2014). One such behavior is avoidance of infected conspecifics; a behavior evolutionarily conserved across metazoans (Curtis, 2014; Kraus et al., 2021). Zebrafish exposed to conspecific alarm substances, thought to be largely composed of skin-commensals released from damaged skin cells, display a fear response characterized by increased darting episodes and erratic behavior which can be learned (Mathuru et al., 2012; Maximino et al., 2018). Interestingly, when adult zebrafish are exposed to heat lysed commensals, there is limited darting response, although there is a fear response like alarm substance-evoked behavior (Chia et al., 2019). Responses to conspecific alarm substance and commensal bacteria are at least in part informed by the olfactory system as evidenced by calcium imaging (Chia et al., 2019; Mathuru et al., 2012). Here adult and larva had decreased swimming velocity 1 d after I.N. IHNV, indicating potential sickness behavior. In support, larvae showed an anxiety-like response to virus by decreasing their speed (Lopez-Luna et al., 2017). Future studies looking at behavioral changes to a variety of nasal pathogens will seek to clarify fear responses and correlate them to neuronal activation states using transgenic, OSN-ablated lines. Further, behavioral changes following multiple olfactory encounters with the same virus is of great interest to us, particularly to understand whether “memory” to olfactory pathogen exposure impacts the brain not only immunologically but also cognitively.

While it is known that viruses can cause damage to the olfactory epithelium and OB in post-viral olfactory dysfunction, there is little information about the immediate responses of OSNs to viruses. Our work has ascertained that OSNs can smell live attenuated IHNV and send action potentials to the OB, measured by EOG (Sepahi et al., 2019). Interestingly, when SARS-CoV-2 S RBD is delivered intranasally, EOG magnitudes in response to food and bile acid odorants decrease significantly (Kraus et al., 2022). While the latter study enforces zebrafish as a model for olfactory dysfunction, the discrepancies between both studies highlight the wide effects that viruses and their proteins can have on olfactory physiology. In the current study, we show through calcium imaging in zebrafish larvae that IHNV specifically activates neurons in the OB.

We did not identify which OSN subsets become activated in response to IHNV, nor did we observe activation of specific glomeruli in the zebrafish OB. This suggests that multiple OSN types may be responsive to IHNV and motivates future experiments comparing different viruses. Furthermore, it is likely that other brain regions are involved in the higher-level processing of these signals into behavioral readouts, and area ripe for further exploration in the context of nasal inflammation and OB responses. The multi-glomerular nature of the response we observed is of particular interest because stimulation with specific odorants has been shown to influence which olfactory receptors neuronal progenitors will express as they mature; this may result in biased expression of olfactory receptors that would alert an organism to danger more rapidly after being around conspecifics shedding pathogen (van der Linden et al., 2020).

Our work shows that neuroimmune responses take place in the OB upon detection of viruses in the olfactory periphery. We provide single cell maps of complex, transient responses, which we hope are useful to many others in the field. One of the challenges of scRNA-Seq is assigning identities to cell clusters not previously characterized, particularly when working with non-mammalian model organisms. For instance, scRNA-Seq of the murine OB has not uncovered many definitive markers for mature OSNs, and few previously suggested markers were found in our zebrafish OB dataset (Tepe et al., 2018; Zeppilli et al., 2021). Therefore, while we captured 5 distinct clusters of neurons, 2 of which represent mature functional subsets and 3 are more progenitor-like by their transcriptional profiles, we cannot determine definitively if we captured interneuron subsets. Our method for capturing single cell suspensions was optimized for neuronal survival, yet neurons only represent about a third of the cells in our dataset after quality control. Our results nevertheless show that detection of viruses in the periphery shapes neuronal transcriptional profiles and populations in the OB. Specifically, at 1 d after I.N. IHNV treatment, neuronal progenitor-like clusters had increased in proportion compared to controls and significantly upregulated expression of genes associated with neuronal differentiation. While acute damage activates quiescent basal stem cells in the olfactory epithelium, chronic damage leads to these horizontal basal cells to enter an inflammatory state (Chen et al., 2017, 2019). Unlike mammals, zebrafish are well known for the regenerative abilities of their adult neurons under inflammatory signaling (Kyritsis et al., 2012). Interestingly, damage to muscle cells can alert distant muscle stem cells into an alert state (Rodgers et al., 2014). In mice, neural stem cells that migrate to the OB and become mature neurons show unique inflammatory profiles as they leave quiescence and can be activated by systemic TNF-α (Belenguer et al., 2021). Here, we present evidence that inflammation in the peripheral OO may prime the progenitor-like neuronal pool in the OB, perhaps as a mechanism to retain plasticity which may be needed if neuronal damage occurs. PACAP is an evolutionarily-ancient neuropeptide, emerging around the protostome-deuterostome divergence, and has a wide variety of functions, including in the olfactory system (Cardoso et al., 2020; Hegg et al., 2003). In our findings, PACAP is exclusively expressed in these progenitor-like clusters. Since PACAP is necessary for survival and differentiation of developing neurons in many model organisms, this is to be expected (Han and Lucero, 2006; Hegg et al., 2003; Irwin et al., 2015). However, there was no direct viral damage in our model because the virus did not reach the OB and therefore the increased populations of PACAP expressing progenitor-like neurons suggests an antiviral role for these cells. Despite the potential anti-viral role of these cells, we did not find that there was higher NFkB expression in the OB following IN viral exposure, indicative that PACAP expression is not simply a result of inflammation but potentially a more specific antiviral program. Indeed, we show that teleost PACAP is directly antiviral in culture (rather than in-vivo), tying neuronal differentiation processes and immune function (Velazquez et al., 2020). These results demonstrate the neuropeptides commonly expressed in the olfactory system can act as neurotransmitters as well as innate immune mediators.

In summary, the present study reveals that virus-elicited olfactory neuronal signals are integrated and translated into behavioral responses in zebrafish. Peripheral viral detection impacts transcriptional programs in neurons, glia and immune cells in the OB creating altering neuronal differentiation programs. Immature neurons in the OB appear to be poised to participate in antiviral immunity via PACAP expression. Our work highlights the importance of investigating neuroimmune communication along the olfactory-brain axis and suggests that brain immunity and behavior are sculpted by chemosensory pathogen detection.

## Materials and Methods

### Zebrafish

Founders of our AB colony were obtained from ZIRC (Oregon, USA) and maintained at the University of New Mexico Castetter Hall Animal Research Care facility’s zebrafish facility at 28°C and a 14-hour light and 10-hour dark cycle. Fish are maintained using a Gemma diet (Skretting). With the exception of larval behavioral and calcium imaging animals used for this study were adults between 6 months and 1.5 years of age. For the behavioral and single cell AB zebrafish were maintained in 6-L mixed sex tanks on the same custom-built recirculating system with solid filtration at 27°C and a 14-hour light and 10-hour dark cycle. For these studies sex was considered as a variable, and data represents evenly both sexes. For larval studies, 5 days post fertilization Tg(vGlut2a:Gal4)^nns20^;Tg(UAS:GCaMP6S)^13a^ zebrafish larvae (Muto et al., 2017; Satou et al., 2013) or Tg(elavl3:GCaMP6S) (Vladimirov et al., 2014) zebrafish larvae were maintained on a Nacre^-/-^ background (White et al., 2008) at the Harvard Neuroscience Department Zebrafish Facility. For larval behavioral studies, 5 dpf wild type AB zebrafish were used as described below. All procedures were done with approval from the University of New Mexico’s Institutional Animal Care and Use Committee under protocol number 22-201239-MC, the University of Nebraska at Omaha protocol number 17-070-09-FC, or the Harvard University protocol number 22-04-5.

### Intranasal treatments of zebrafish

For intranasal treatments animals, either AB strain zebrafish or Tg(NFkB:EGFP*)^nc1^* (Kanter et al., 2011) were anesthetized with 0.04 mg/ml neutral buffered Tricaine (Syndel) in tank water for 1 m and then transferred to an absorbent boat soaked with anesthesia solution for the duration of treatment. If endpoints were the next day, animals were moved to the system overnight. Live attenuated infectious hematopoietic necrosis virus (IHNV) was at 2×10^8^ plaque forming units (pfus) as previously described (Fryer et al., 1976; La Patra et al., 2004) and approximately 5µl per nare was delivered using a microloader pipette tip (Eppendorf, 930001007). Animals were then moved to a recovery tank and monitored until end point time. Sterile PBS (approximately 5µl per nare) was a control for the anesthesia and mechanical perturbation of intranasal treatments.

### qPCR detection of virus and RT-qPCR for chemokine expression

Olfactory organs and OB from 3 individual animals were dissected out and separately pooled in 500μl of Trizol (Cat 15596026), then snap frozen. Tissue was homogenized by adding two tungsten beads to thawed trizol and shaken at 30 Hz for 5 m. RNA was extracted following the Trizol protocol as previously described for this tissue (Kraus et al., 2022). RNA was quantified and standardized for cDNA synthesis using Superscript III first stand system (Thermofisher 18080051). qPCR was performed using SSOadvanced supermix (Biorad 1725270) at a T_M_ of 62°C. Primers for N protein of IHNV are F: 5’-GGTCGCCGAACTTCTGGAA-3’ and R: 5’-GTAGGGCGCAGGTGAAGAGG-3’.

### Live imaging of neuronal calcium signaling in zebrafish larvae

Five days post-fertilization Tg(vGlut2a:Gal4)^nns20^;Tg(UAS:GCaMP6S)^13a^ zebrafish larvae (Muto et al., 2017; Satou et al., 2013) on a Nacre-/- background (N =5) were embedded in 2% low melting point agarose and covered with facility water. Additionally, pilot studies were conducted on 5 dpf Tg(elavl3:GCaMP6S) larva (Vladimirov et al., 2014). The area from the olfactory pits to the most rostral part of the animal was freed from agarose using a scalpel. The tail was also freed by removing agarose posterior to the swim bladder. Heart rate was monitored to ensure health. 2-P imaging of the olfactory epithelia and the olfactory bulb were done as previously described using random 10 s pulses of odorants directed to the olfactory pits every 2 minutes (Herrera et al., 2021). Odorants were either live attenuated IHNV, DMEM medium + 10% FBS as a negative control, or 100 µM alanine as a positive odorant control for the duration of the experiment. Image acquisition was performed as done previously at 0.7 Hz (Herrera et al., 2021). Each plane was imaged for 15 minutes, with up to 10 planes imaged per fish.

To identify cue-specific cells, we generated stimulus-based regressors by convolving the stimulus presentations with the GCaMP fluorescence kernel. We then determined cells to be active as those whose best fit to one of these regressors fell into the top 10% of fits across all regressors in a given fish. When then calculated the proportion of cells that belong to each possible response type as the fraction of those top 10% of cells with a top fit to that category’s regressor. Data analyses of calcium signals were performed as described.

### Larval behavioral studies

All experiments took place between 10am and 6pm. 6 dpf larvae, chosen for their fully developed olfactory system and transparency (Li et al., 2005) were placed in the setup 5 min before the experiment started. In a custom-built setup, the fish experienced a constant water flow (about 34 ml/min) in a 4 cm x 4 cm chamber. Water was constantly flowing in on one side of the chamber and pumped out at the opposite side of the chamber. Meshes on both ends of the chamber prevented the fish from getting into the pump if they stopped swimming. A custom written software controls the system that allows for automatic stimulation with any liquid, for example a virus diluted in water. Two arduino controlled valves are switched between a constant water flow and the e.g., virus that is added to the constant water stream. Every fish received three 10-second-long presentations of either the virus or a vehicle and in addition three 10-second-long presentations of water as a control for valve switching related changes in behavior. The order of the presentation of virus and control was alternated between fish. The inter stimulus interval between the presentations was 150 seconds. The virus/vehicle was cleared from the chamber at 120 seconds post-delivery. The timing of exposure and clearance was determined based on the flow of the chamber. We calculated that it takes 30s for the virus or vehicle to reach the fish and another 120s to exit the chamber, resulting in 150s.

Fish were filmed from above using FLIR pointgrey grasshopper IR camera at 5 frames per second. Fish tracking was done using a costume software written in Matlab (for full details see Harpaz et al., 2018, PNAS). Briefly, we created a background model by calculating the median pixel intensity of 500 equally spaced frames along the movie. We then subtracted this model from all frames along the movie and thresholded the subtracted image to detect the fish as the darkest blobs in the image. Finally, we took the center of mass of the blobs as the fish coordinates 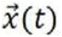.

Preprocessing and filtering of data was performed as follows. Fish coordinates 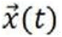 were smoothed using a Savitzky-Golay filter of ∼1/3 second (10 frames). Velocity was calculated as, 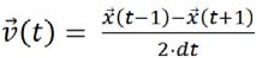, and the speed is given by the magnitude of the velocity 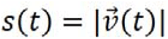. To remove spurious data points that are likely due to tracking errors, we excluded all data points with speed equal to zero and points where the time difference *dt*, between two consecutive tracked coordinates was larger than 0.04s, which indicated a temporary loss of the fish by the tracking software.

We estimated the strength of the water flow dragging the fish back (see ‘experimental setup’) by first finding all instances where the fish was facing in the direction of the upcoming water flow with heading angle θ between 90° and −90° (swimming directedly towards the flow is defined as 0°) yet was moving ‘backwards’ with the flow. We then averaged all the fish velocity components that were parallel to the flow direction 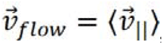, where 〈…〉 represent average over all velocity measurements.

Finally, we calculated the corrected velocity vector of the fish, correcting for the fact that the fish is swimming against the constant water flow 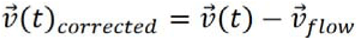, and accordingly we have 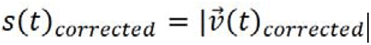. We note that this calculation assumes a homogenous flow across the arena and therefore is only a representation of the average flow experienced by the fish.

### Adult behavioral studies

Behavioral assays were conducted as previously reported (Baker et al., 2018). In short, square plexiglass arenas with an inner width of 12 inches were filled with 4 L of system water and left to settle inside a white curtained area. Camera was mounted above, and focused. Fish were alternatively intranasally treated with media (DMEM+5% FBS, used to grow IHNV) or live attenuated IHNV. Animals were anesthetized in 0.04 mg/ml Tricaine (Syndel) for 1 m, intranasally treated, then left in a recovery tank for 5 m. Then they were placed in the arena and allowed to settle for 30 seconds before video recording for 5 m. Arenas were washed between trials. No water was allowed to cause color discrepancies under the arenas. Analysis of behavior was performed with EthoVision software. Animals that traveled less than 10 cm were removed from analysis.

### Single cell isolation and capture

After animals were euthanized, the OBs were dissected out and immediately placed in individual tubes with Neurobasal medium (Gibco Cat# 21103049) supplemented with 0.04% BSA and 5% FBS. This supplemented media was used throughout cell isolation and capture. OBs were gently rocked for 45 m, then all (N = 6) samples for a single treatment were mashed through a 100mm cell strainer into 3ml of supplemented media. Cells were washed 2 times by spinning 400 x g for 8 m, carefully removing supernatant and resuspending in fresh medium. When cells were pipetted up and down 3 seconds were counted for plunger movements up and down. Rainin low bind tips were used throughout. Cells were then sorted for viability using propidium iodide. Cells were washed once and resuspended in 60 ml. Ten ml cells were used for counting with a hemocytometer and 10,000 cells were loaded in 40ml to the 10x chromium cell capture machine after running through a 40µm Flowmi cell strainer (Bel-art catalog no. H13680-0040) to remove clumped cells. Libraries were prepped and sequenced by University of Nebraska DNA Sequencing Core. Sequencing metrics can be found in Figure S3A.

### Analysis of single cell RNA-Seq

Fastqs were generated and processed into count matrices using Cell Ranger v3.0 under default settings using a reference generated using the GRCz11 zebrafish genome. In R (v3.4.1) and Surat (v3.1.1), matrices were imported into a Seurat object for analysis. As quality control, cells with less than 200 or greater than 2500 features and greater than 5% of features belonging to the mitochondria were removed. Counts were then normalized using the ‘LogNormalize’ method and a scale factor of 10,000. Using FindVariableFeatures under the ‘vst’ method, 2000 variable genes were found. Treatments were then integrated using the CCA method and scaled using default settings (Butler et al., 2018). Dimensional reduction was done first with PCA analysis using 30 significant principal components determined using Jackstraw and Elbow plot analysis. Secondary dimensional reduction was done with tSNE. Clusters were found using ‘FindNeighbors’ and ‘FindClusters’ sequentially at a resolution of 0.5 and 30 significant PCs. Differential expression was done using ‘FindMarkers’ set to default between treatments within each cluster and significant features with Bonferroni adjusted p-values of less than 0.05 were then exported for gene ontology. Plots were made in R using a combination of Seurat and ggplot2 (v3.3.5) elements. Gene ontology was done using Metascape, which produced the biological process bar plots (Zhou et al., 2019).

### Antiviral and cell viability assays

EPC cells were maintained at 15 in MEM (Corning) supplemented with 10% FBS. IHNV stocks were grown in EPC cells as previously described (Ma et al., 2019). The cytotoxicity of the tested extract was evaluated using MTT assay (CyQUANT™ MTT Cell Viability Assay). EPC cells (10000 cells/well) were prepared in 96-well plates for 24h. Cells were exposed to concentrations of PACAP-85% or amidated PACAP-96% samples in quadruplicate (10µM, 1µM, 100nM, 10nM and MEM vehicle control), followed by 48h incubation at 15 °C. Cell viability was measured using the CyQUANT MTT cell viability assay (Thermo Fisher Scientific Cat #V13154) following the manufacturer’s protocol.

Three different assays were applied to investigate the antiviral activity of PACAP with various concentrations: pretreatment, competition, and post-treatment. All assays were performed in quadruplicate. For amidated PACAP-96%, concentrations of 1µM, 800nM, 600nM, 400nM and 200nM were used; for PACAP-85%, 50µM, 40µM, 30µM, 20µM and 10µM were used.For all treatments, EPC cells (10^6^ cells/well) were prepared in 24-well plates for 24h. All infections were performed at a Multiplicity of Infection of 0.05. In the pretreatment assay, cells were treated with PACAP-85% or amidated PACAP-96% for 2h, then washed before infection with IHNV for 1h. In the competition assay, PACAP-85% or amidated PACAP-96% and IHNV (10^5^ pfu/ml) were mixed and incubated at 15 °C for 2h. Mixtures were added to the monolayers of EPC cells for 1h. In the post-infection treatment assay, the monolayers of EPC cells were incubated with IHNV for 1h, rinsed, then treated with PACAP-85% or amidated PACAP-96% for 2h. Cells were washed twice with MEM, then incubated in MEM with 2% FBS at 15 °C and observed with an inverted microscope. After 7 days incubation, cell viability was measured using the CyQUANT MTT cell viability assay (Thermo Fisher Scientific) following the manufacturer’s protocol. Virus-infected EPC cells and non-treated cells served as infection control and blank control, respectively. The relative survival of cells was calculated as: [(OD value of treatment group − OD_570_ value of infection control)/(OD_570_ value of blank control − OD_570_ value of infection control)] × 100.

### Immunofluorescence staining

30 µm cryosections were stained with a polyclonal anti-PACAP-38 antibody (Abcam, ab216627). In short, heads were fixed overnight in 4% methanol-free PFA at room temperature, followed by 5 days in 10% EDTA, pH 7.2. Samples were cryoprotected in 33% w/v sucrose in ddH2O until saturation, followed by flash freezing in Tissue-Tek OCT Compound (Sakura Finetek USA, catalog no. 4583). Samples were processed into 30 µm sections with a Leica CM3050S (Leica, Wetzlar, Germany) onto Superfrost Plus Slides (VWR catalog no. 48311-703). OCT was removed from sections displaying the OB by 3 rinses for 5 m each in 1x PHEM, then permeabilized for 10 m with 0.5% Triton X-100 in 1x PHEM, followed by 2 rinses for 5 m each in 1x PHEM. For anti-PACAP staining, antigen retrieval was performed by covering samples in citrate buffer (0.1 M, pH 6.0) for 5 m at 90°C, followed by 3 rinses for 5 m each in 1x PHEM. Samples were blocked for 1 hours with BlockAid blocking solution (Thermo Fisher Scientific, catalog no. B10710)). Samples were incubated overnight at 4°C with anti-PACAP-38 antibody (Abcam, catalog no. ab216627, 1:100) or normal Rabbit IgG (Santa Cruz Biotechnology catalog no. sc-3888) in Pierce Immunostain Enhancer (Thermo Fisher Scientific catalog no. 46644) with 0.1% v/v Tween and 1% w/v Bovine Serum Albumin. Following incubation with primary antibodies, samples were rinsed twice for 5 m each in 1x PHEM. Samples were then incubated for 2 h with AlexaFluor 647 Goat anti-Rabbit (1:300, Jackson Immunoresearch catalog no. 111-605-144) diluted in 1x PHEM with 0.1% v/v Tween 20, 1% w/v Bovine Serum Albumin, 10% v/v Normal Donkey Serum, and 200 mg/ml DAPI. Samples were then rinsed three times for 10 m each, then mounted with Prolong Glass Antifade Mountant with NucBlue (Thermo Fisher Scientific catalog no. P36981). For each round of staining, an isotype control of normal Rabbit IgG was included. For Edu detection, zebrafish were injected intraperitoneally 1 day before sacrifice as previously described (Lindsey et al., 2018) and detected following the click-it Plus for imaging protocol (Thermo Fisher catalog no. C10640). Confirmation by adsorption of anti-PACAP-38 with recombinant PACAP-38 protein was performed by incubating the antibody with ten times molar concentration of the recombinant protein for 90 m at room temperature prior to immunostaining. For anti-EGFP staining of *Tg(NFkB:EGFP)* fish, the protocol was followed as above with the exception of the antigen retrieval step. Specifically, chicken IgY anti-EGFP (1:500, Aves Labs catalog no. GFP-1010) was used as primary antibody followed by secondary antibody staining using Cy5-labelled donkey anti-Chicken (1:300, Jackson Immunoresearch catalog no. 703-175-15). Samples were imaged on a Zeiss LSM 780, equipped with an Xcite halogen lamp. All images for quantification to be compared with each other were imaged on identical settings. Images were captured as a .czi file, which was imported into Fiji using the Bioformats importer. ROIs were placed around the center region of the OB to exclude the background edge and mean gray value intensity measurements of each ROI were taken.

### Statistical analysis

To check that data was normally distributed, an F test, Bartlett’s test, or D’Aggostino test was done on all data. Parametric data with 2 groups for comparison were analyzed using a Student’s T test or a Wilcoxon-signed rank test for comparison within groups. Population proportion analysis of cells in each cluster between clusters was analyzed using ‘prop.test’ and p-values were adjusted using ‘p.adjust’ set to ‘bonferroni’ method. All analyses were performed in prism (v 9.3.1). For larval behavioral studies, statistics were performed as previously described (Herrera et al., 2021). Odorant-selective rois were identified by selecting ROIs with activity during the 30s following the onset of the chosen stimulus that was 1.6 times greater than the sum of the activities during the other two stimuli over the same time.

## Data availability

NCBI Bioproject Number is PRJNA1520453.

## Acknowledgements

This work was partly funded by the ADVANCE UNM Women in Science Award to IS and the National Science Foundation award # 1755348. AK received support from the UNM Center for Advanced Research Computing (CARC) and BJG was funded by the HHMI Gilliam fellowship. The authors would like to thank The University of Nebraska DNA Sequencing Core receives partial support from the National Institute for General Medical Science (NIGMS) INBRE - P20GM103427-14 and COBRE - 1P30GM110768-01 grants as well as The Fred & Pamela Buffett Cancer Center Support Grant - P30CA036727. KJH, HZ, RH, and FE received support from National Institute of Neurological Disorders and Stroke U19-NS104653-03. We thank Drs. Yamila Carpio, Mario Pablo Estrada and Brian Dixon for sharing the recombinant PACAP-38. We thank Dr. Michael Paffett and the New Mexico Comprehensive Cancer Center’s Fluorescence Microscopy and Cell Imaging core for technical support and Drs. Lijing Bu and Marijan Posavi for bioinformatics support. This publication’s contents are the sole responsibility of the authors and do not necessarily represent the official views of the NIH, NINDS, or NIGMS.

---

**Supplemental Figure 1:**
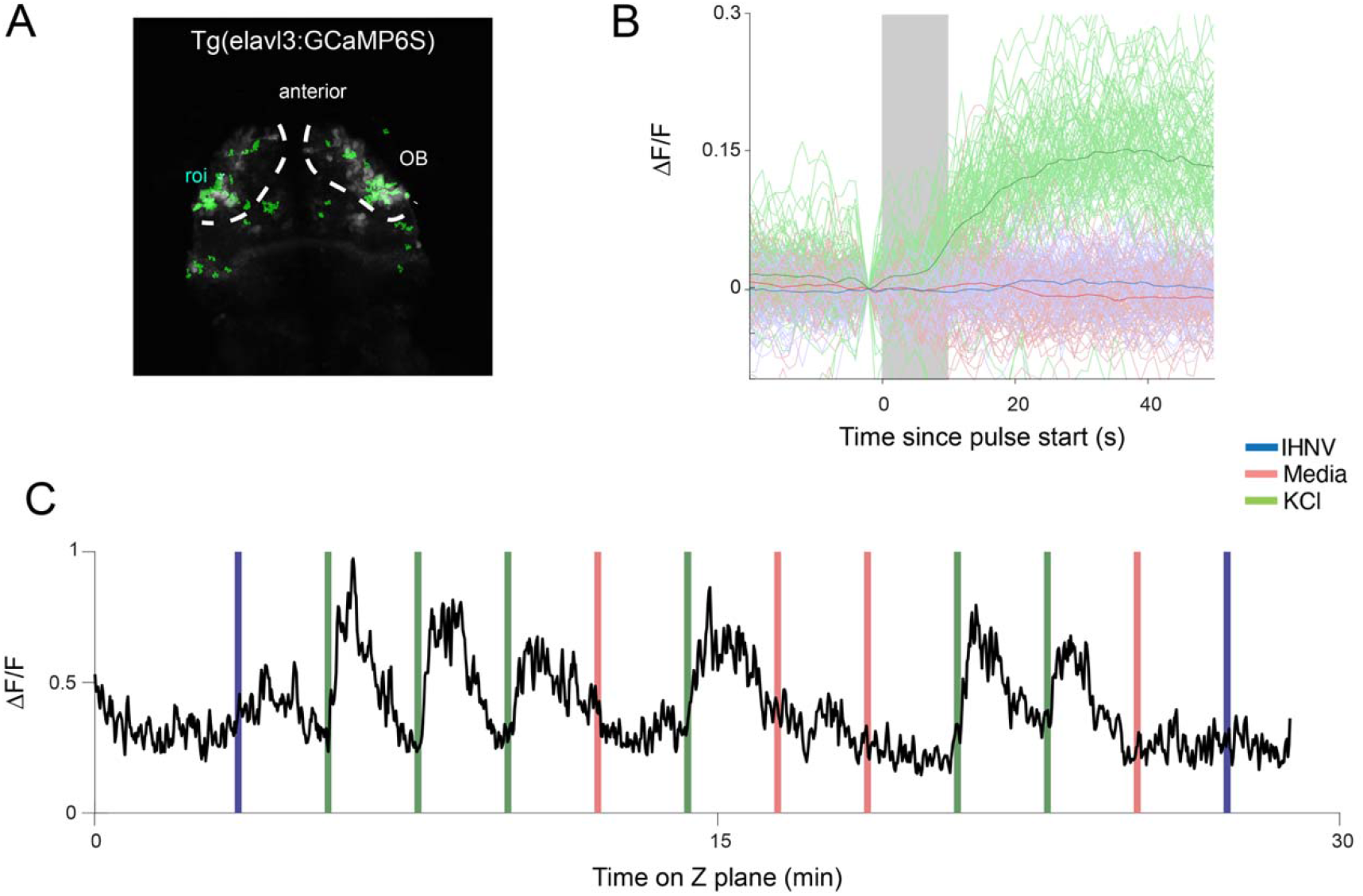
(A) Olfactory system of a Tg(elavl3:GCaMP6S) larva imaged by 2-P microscopy. Odorants were applied to the olfactory pit randomly every 2 m and animals imaged for 6 h., OB = olfactory bulb. Indicated in green are ROIs selective to KCl that do not respond to the other stimuli. (B) Stimulus triggered average changes in fluorescence (F/F_0_) following each stimulus for cells that show neuronal activation specific to the positive (KCl) control (green line) but not the IHNV (red line) virus or negative control (Media, blue line). Mean response across KCl-selective cells. (C) F/F_0_ traces of the highlighted ROI in A. Vertical lines color coding indicates the type of stimuli delivered at each interval.

**Supplemental Figure 2.**
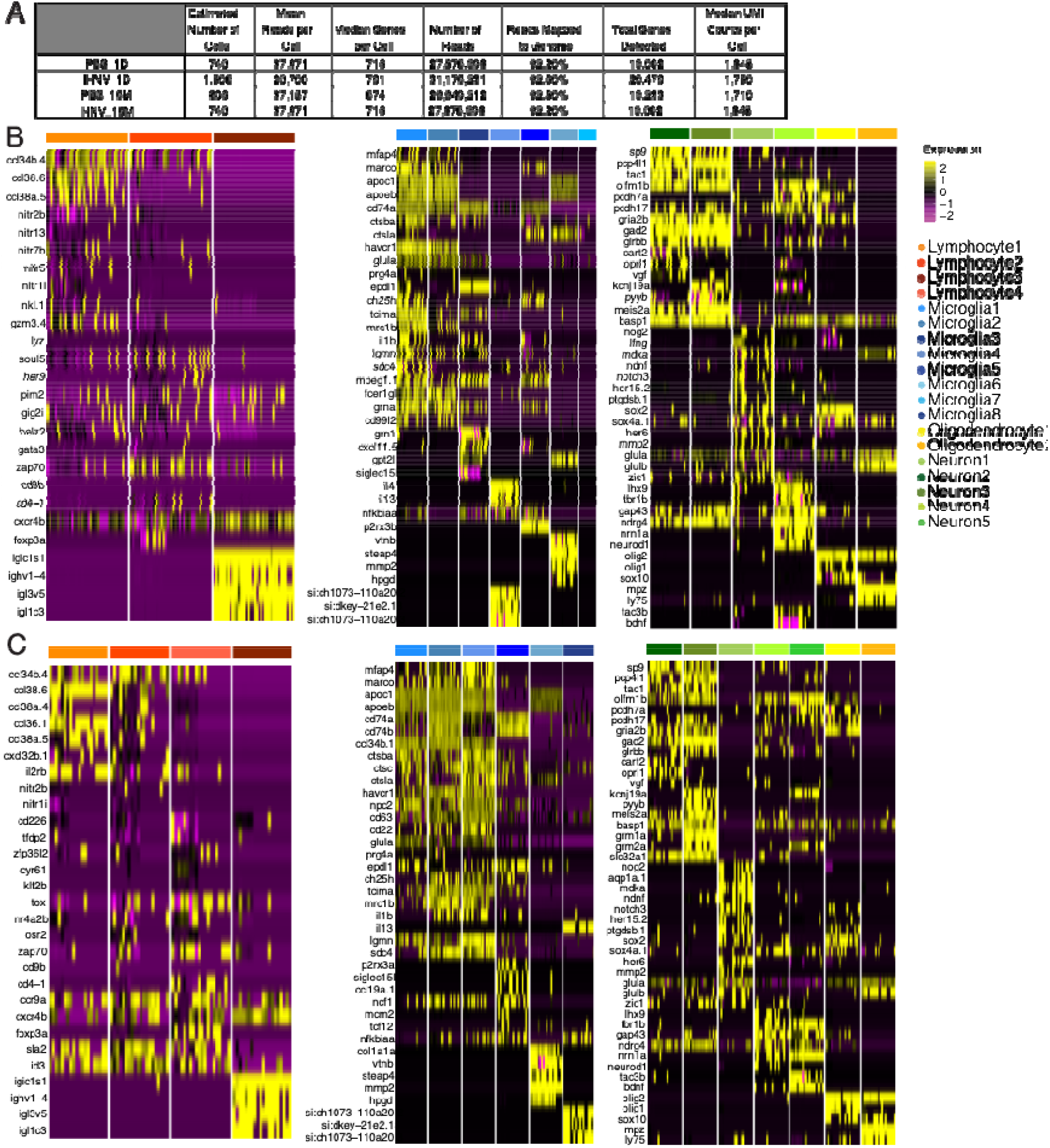
Transcriptional landscape of the adult zebrafish OB 15 m and 1 d after intranasal viral delivery. (A) Metrics from each sample collected by 10x Genomic Chromium platform after sequencing on Illumina MiSeq. (B) Heat map of cluster markers for lymphocyte, microglial, and neuronal clusters from CCA integrated Vehicle and IHNV treated cells from the OB 15 m after intranasal treatment. (C) Heat map of cluster markers for lymphocyte, microglial, and neuronal clusters from CCA integrated Vehicle and IHNV treated cells from the OB 1 d after intranasal treatment.

**Supplemental Figure 3.**
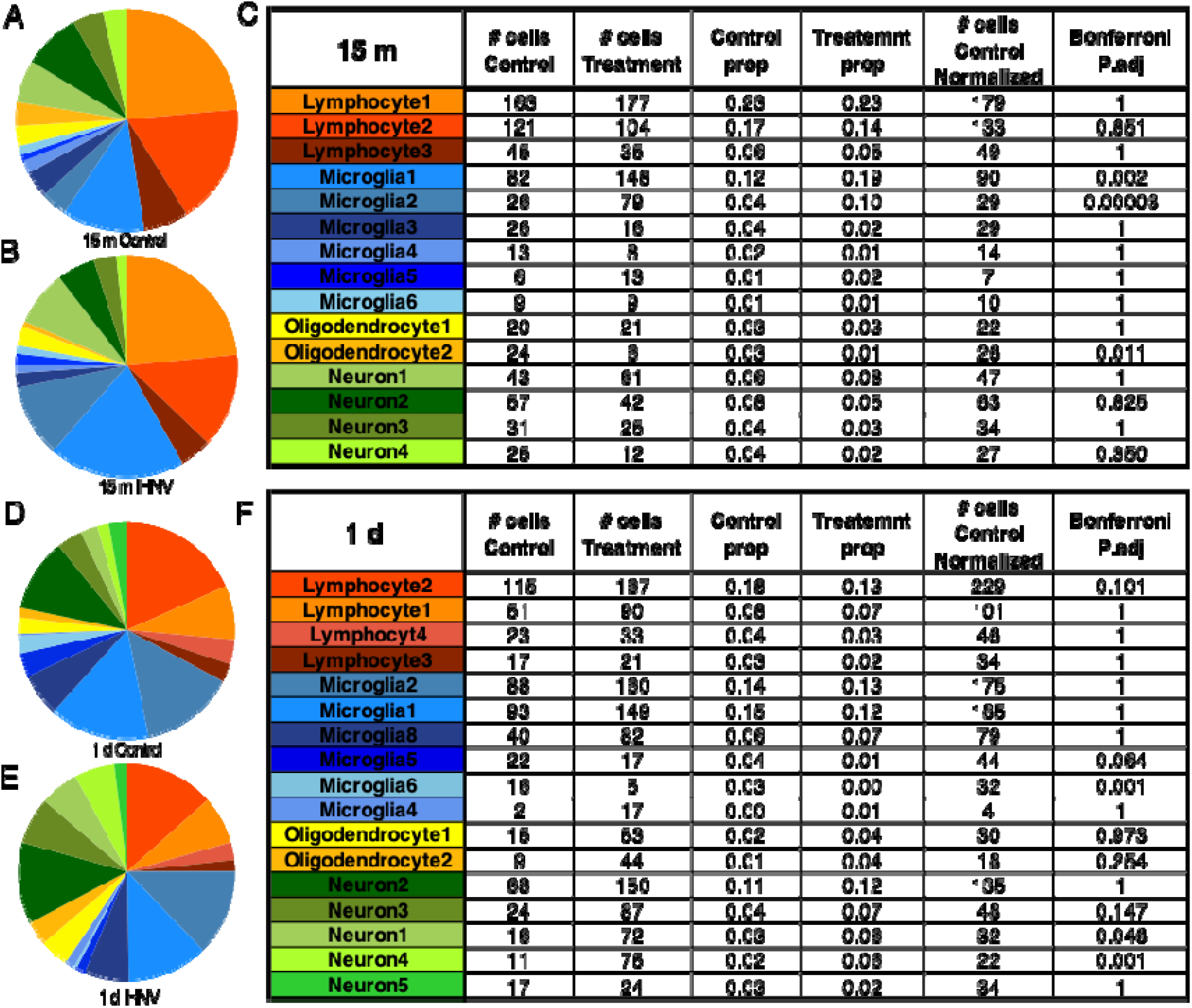
Cellular landscape of the adult zebrafish OB 15 m and 1 d after intranasal delivery of vehicle (PBS) and virus. (A&B) proportional analysis of cells in each cluster in each treatment for 15 m with statistical analysis in (C). (D&E) proportions analysis of cells in each cluster in each treatment for 15 m with statistical analysis in (F) When the *P*-value<0.05, the null hypothesis that population sizes are the same in both treatment groups i rejected.

**Supplemental Figure 4.**
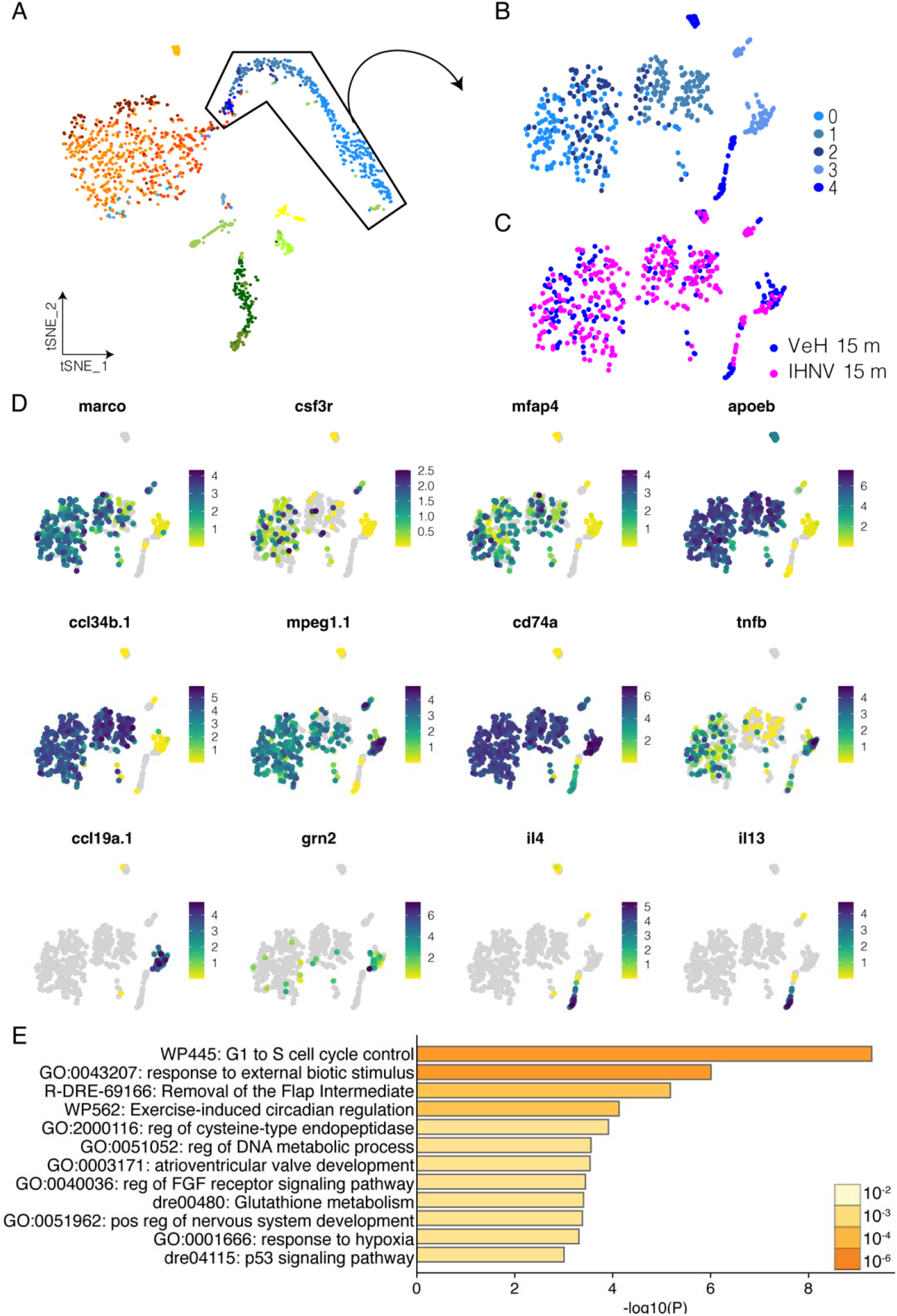
Microglia and macrophage populations in the adult zebrafish OB are diverse. (A) tSNE of the whole OB 15 m after I.N. IHNV to represent the microglial cell types that were subclustered in (B) and represented as either vehicle (VeH) control (blue) or IHNV treated (pink) (C). (D) Feature plots showing markers of microglia (*marco, csf3r, mfap, apoeb)* and ameboid like microglia (*ccl34b.1)* as well as macrophages (*mpeg1.1, ccl19a.1, grn2).* (E) Gene ontology of differentially regulated genes from microglial clusters 15 m after I.N. IHNV compared to vehicle. Gene ontology was run on Metascape (Zhou et al., 2019).

**Supplemental Figure 5.**
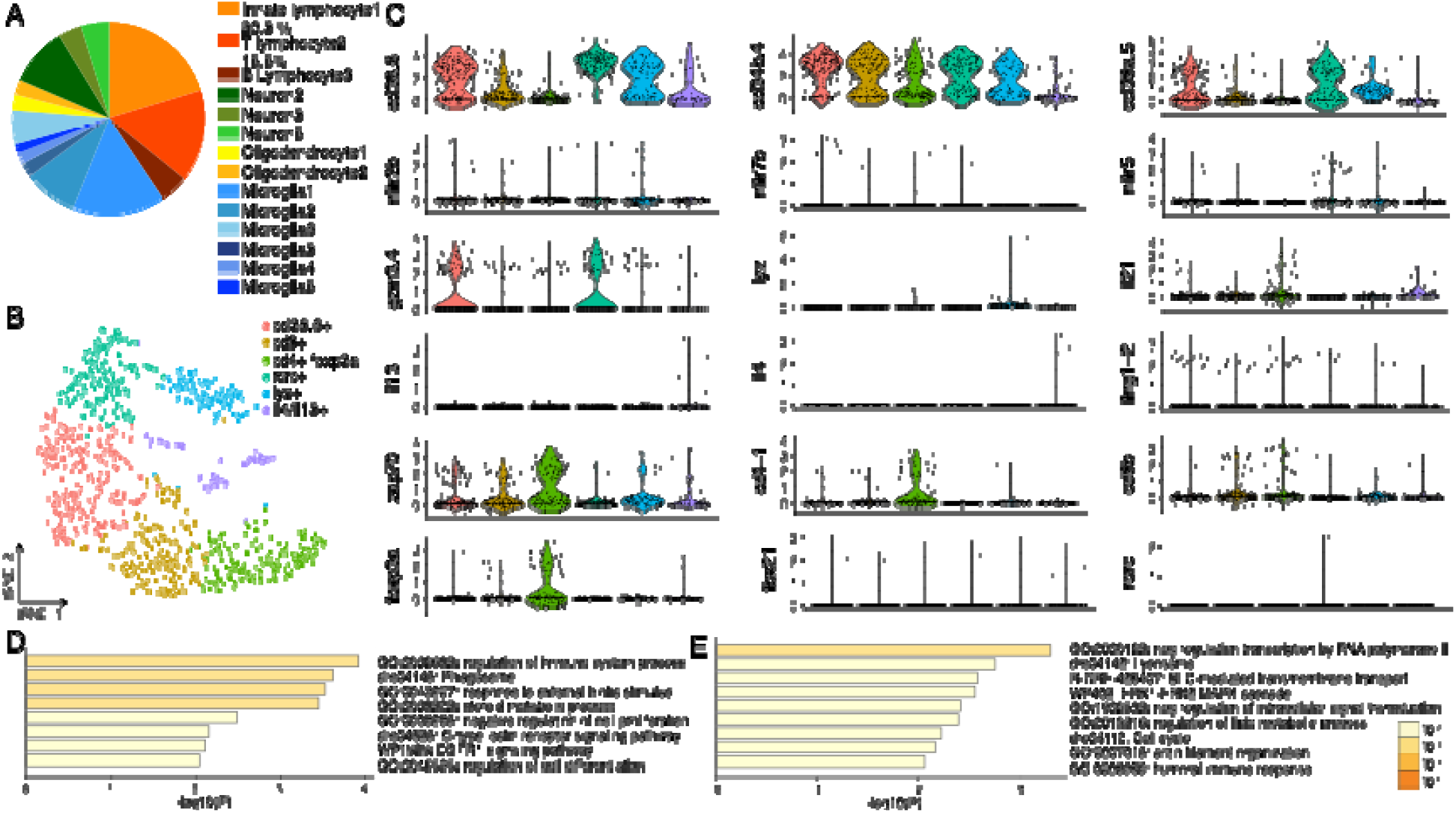
Diverse lymphocyte clusters are found in the adult zebrafish OB. (A) Proportions of clusters from figure 6 of only control cells shows innate-like lymphocytes are 20% of cells captured from the control OB and T lymphocytes2 are 15% of cells captured from the control OB. (B) t-SNE of innate/T lymphocyte subsets from the OB show 6 unique phenotypes. (C) Violin plots of markers of natural killer (*ccl38.6, ccl34b.4, ccl38a.5, nitr*), cytotoxic (*gzm3.4, lyz, ifng1-2, tbx21*), type II innate-like lymphocytes (*il13, il4*), type III innate like lymphocytes (*il2, rorc*) and Tregs (*foxp3a, cd4)*. (D) Gene ontology of significantly differentially expressed genes from lymphocytes in the OB 15 m after I.N. IHNV compared to 1d I.N. vehicle treated lymphocytes. (E) Gene ontology of significantly differentially expressed genes from lymphocytes in the OB 1 d after I.N. IHNV compared to 1d I.N. vehicle treated lymphocytes. Gene ontology was run on Metascape (Zhou et al., 2019)

**Supplemental Figure 6.**
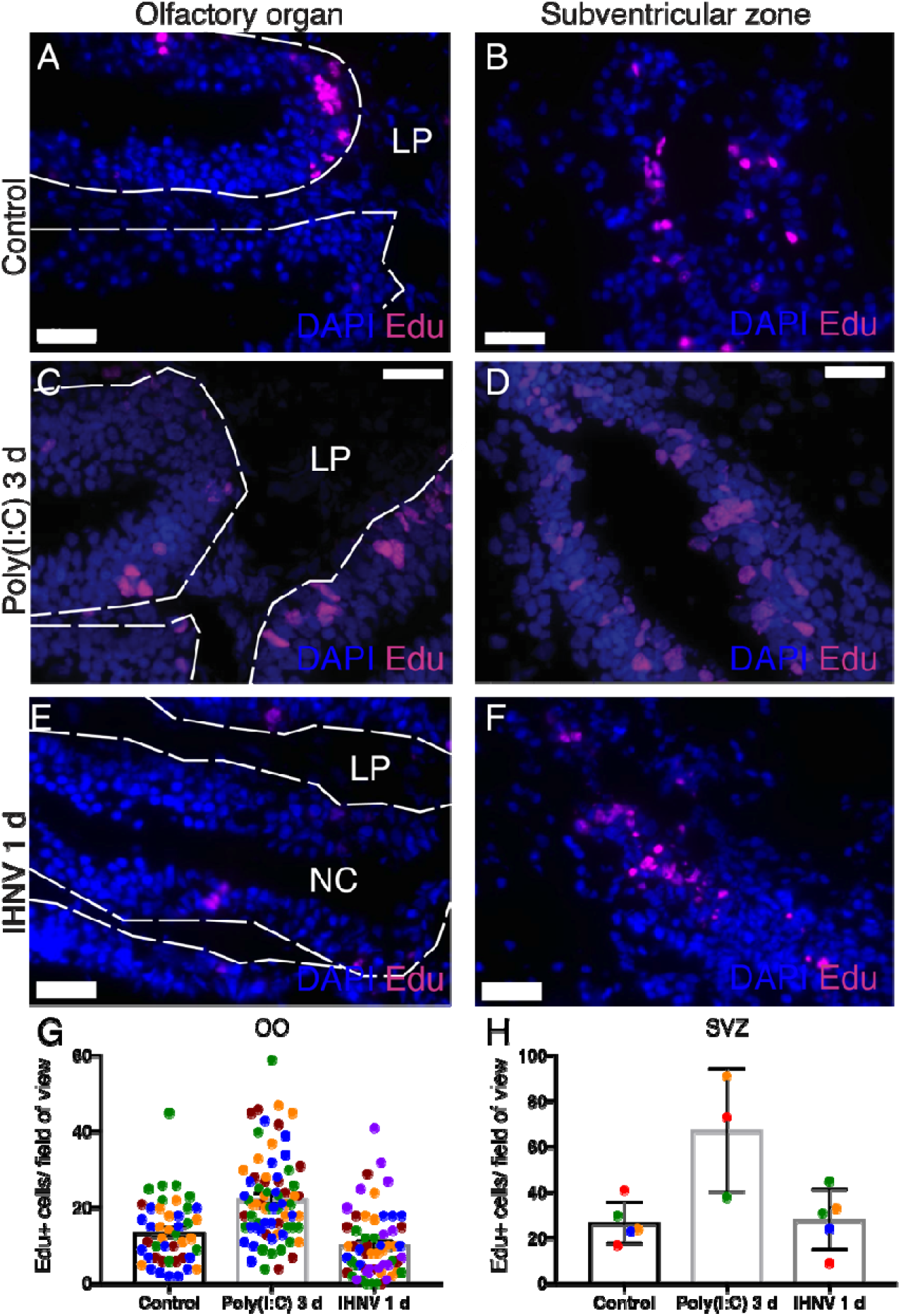
I.N. IHNV does not induce proliferation in the adult zebrafish olfactory organ or the subventricular zone of the telencephalon. Proliferation detected using Edu (pink) in control (A) 3 d I.N. Poly(I:C) (C) or 1 d I.N. IHNV (E) in the olfactory organ (OO) quantified in (G). Proliferation detected using Edu in control (B) 3 d I.N. Poly(I:C) (D) or 1 d I.N. IHNV (F) in the subventricular zone (SVZ) quantified in (H). scale = 20mm. N= 3-5. Each color indicates a separate animal and each dot a field of view at 60x. Statistics are Student’s T test between control and each treatment and not significant for the IHNV treatment.

**Supplemental Figure 7:**
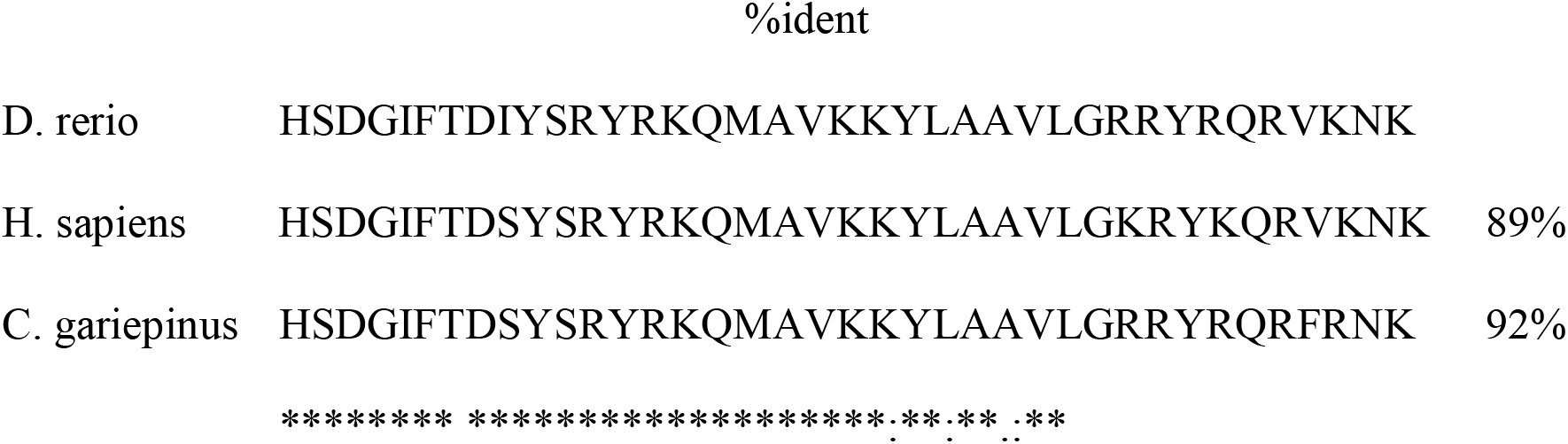
Alignment of PACAP-38 sequences from *D. rerio* (NP_001259010.1:96-133), *Homo sapiens* mature PACAP-38 peptide (NP_001093203.1:132-169), and *C. gariepinus* from (Semple et al., 2019). Percent identity is compared to *D. rerio*.

**Supplemental Figure 8:**
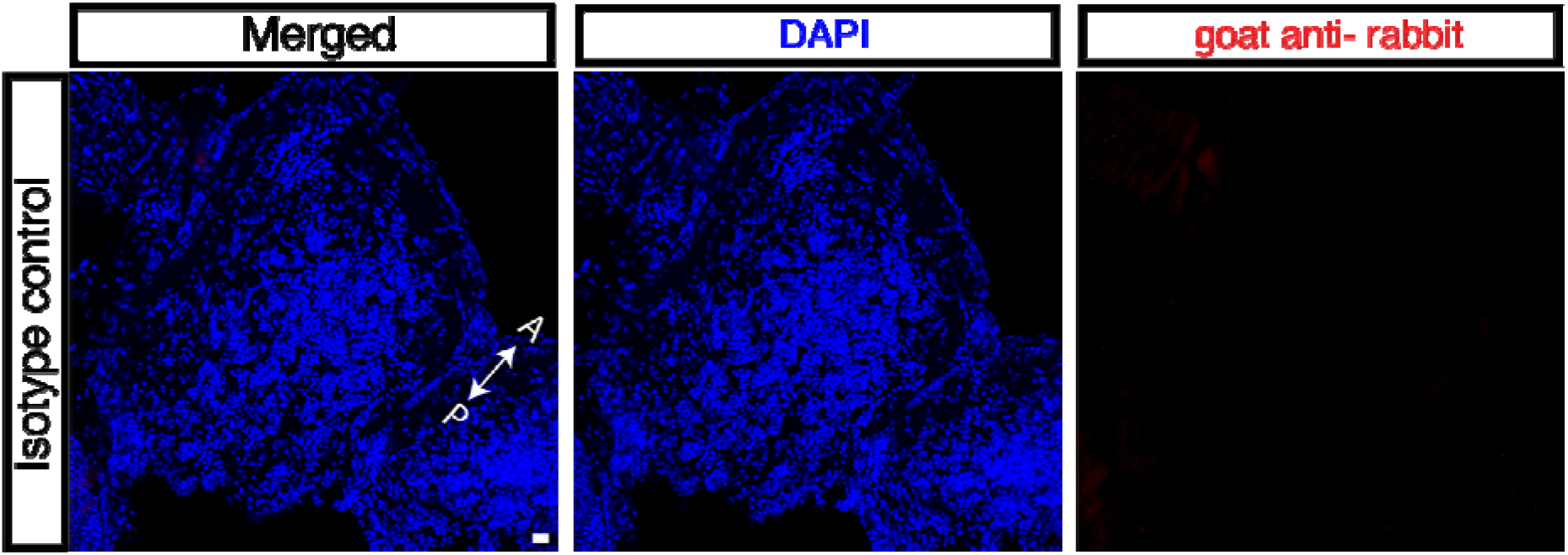
Isotype control staining of coronal adult zebrafish OB cryosections for anti-PACAP-38 staining. Scale bar = 20um.

**Supplemental Figure 9:**
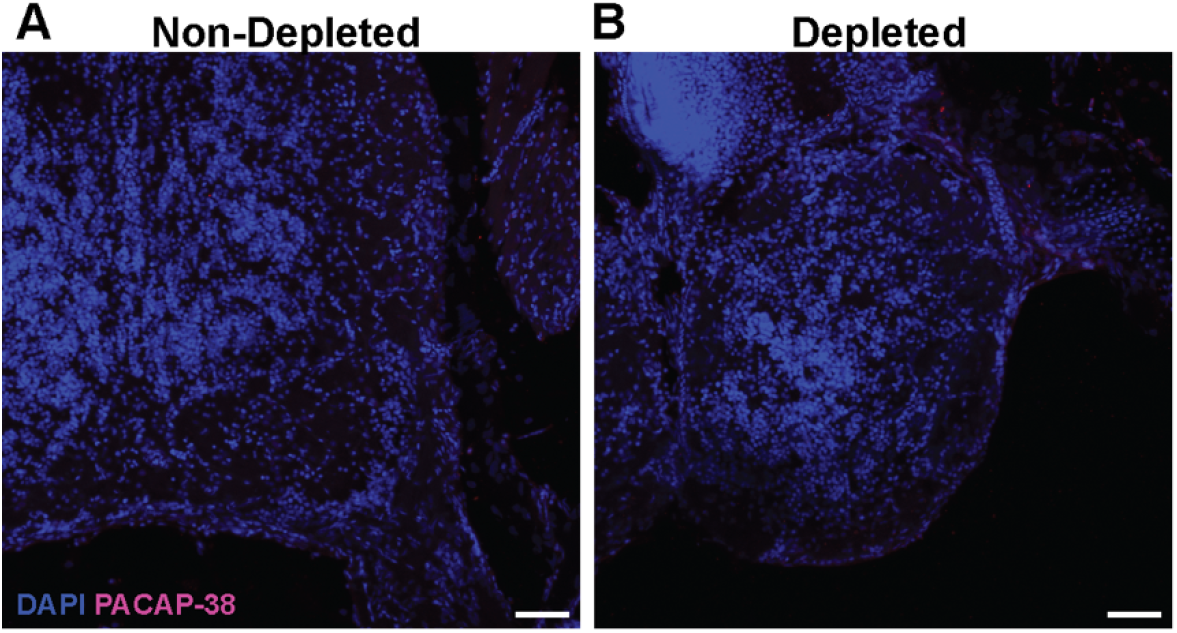
PACAP staining of the adult zebrafish OB following antibody blocking with recombinant PACAP-38. (A) Immunofluorescence image of a control OB stained with anti-PACAP-38 antibody. (B) Immunofluorescence image of a control OB stained with anti-PACAP-38 antibody previously blocked with recombinant fish PACAP-38 showing that signal is lost. Scale bar: 50μm

**Supplemental Figure 10:**
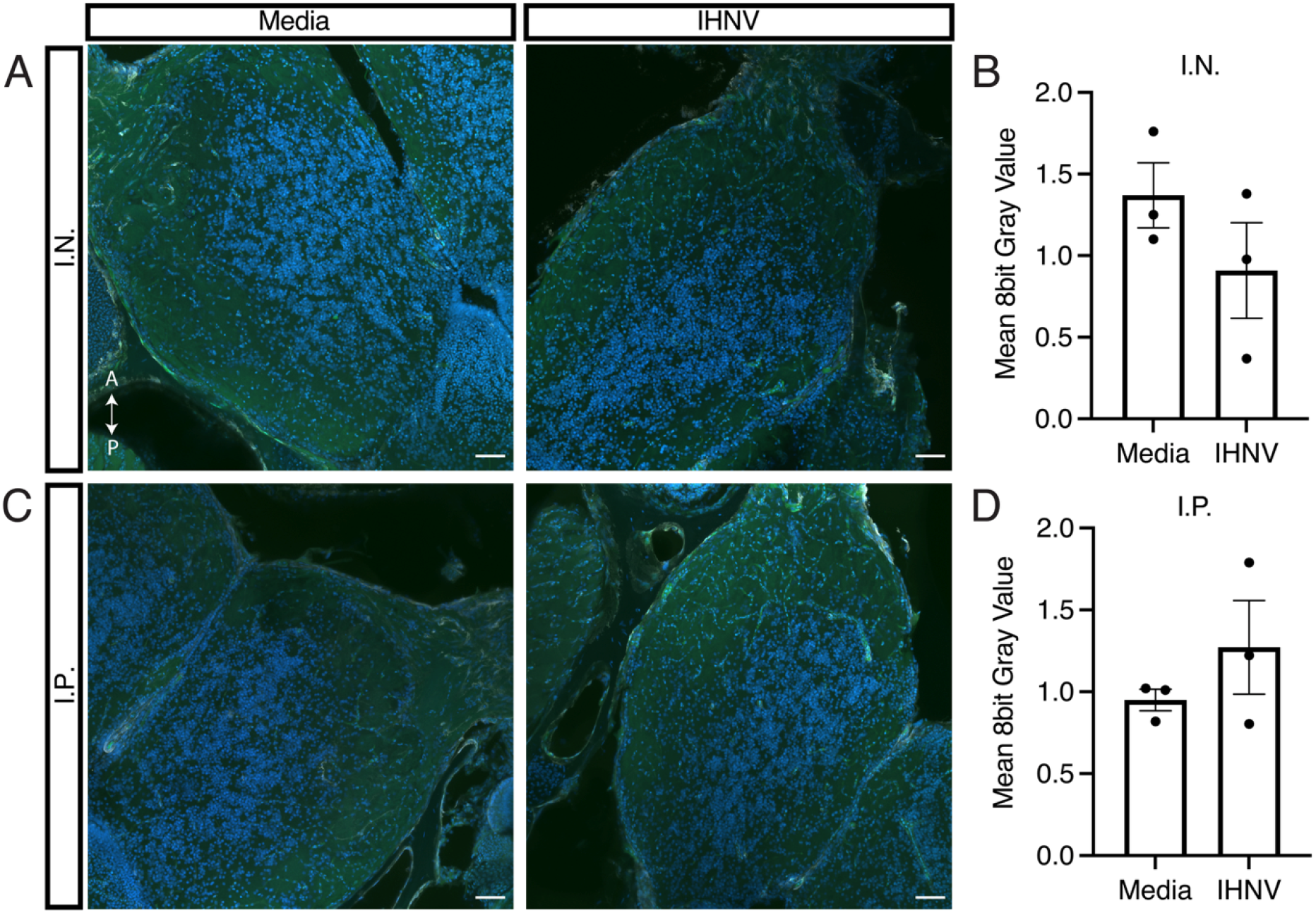
NFκB expression in the adult AB zebrafish OB following IN and IP administration of IHNV. Images of anti-EGFP staining of the of *Tg(NFkB:EGFP)* transgenic fish which received media or IHNV intranasally. (B) Quantifications of images in (A). OB images of fish which received IP media or IHNV. Quantification of (C). Statistics were performed as a Student’s T test. Scale bar: 50μm. IN, intranasal; IP intraperitoneal.

**Supplemental Figure 11:**
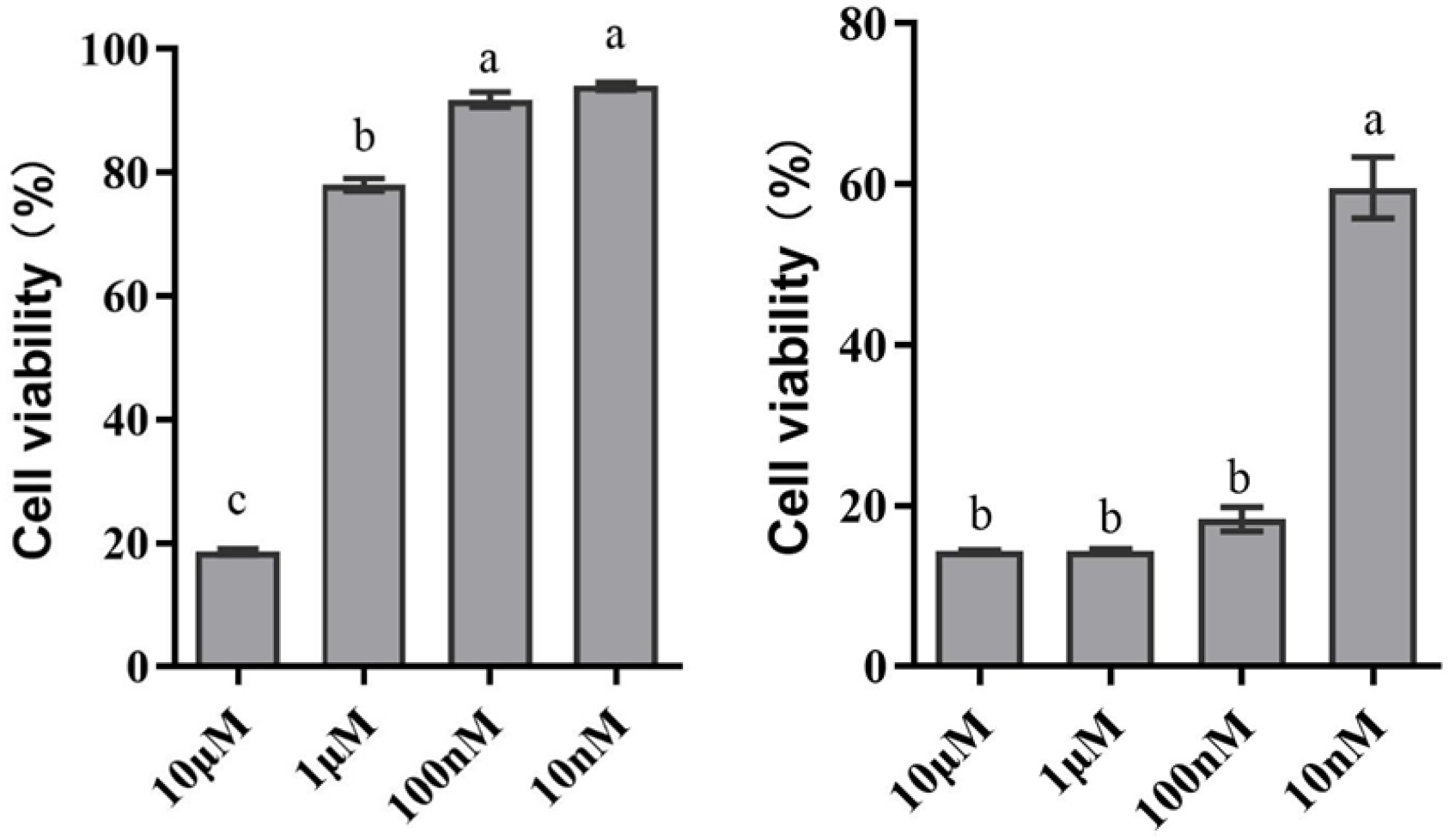
Cytotoxicity of PACAP in EPC cells. (Left, amidated PACAP-96%; right, PACAP-85%;). Different letters indicate statistically significant differences by one-way ANOVA. Results are indicative of two independent experiments.

**Supplemental Table 1.**
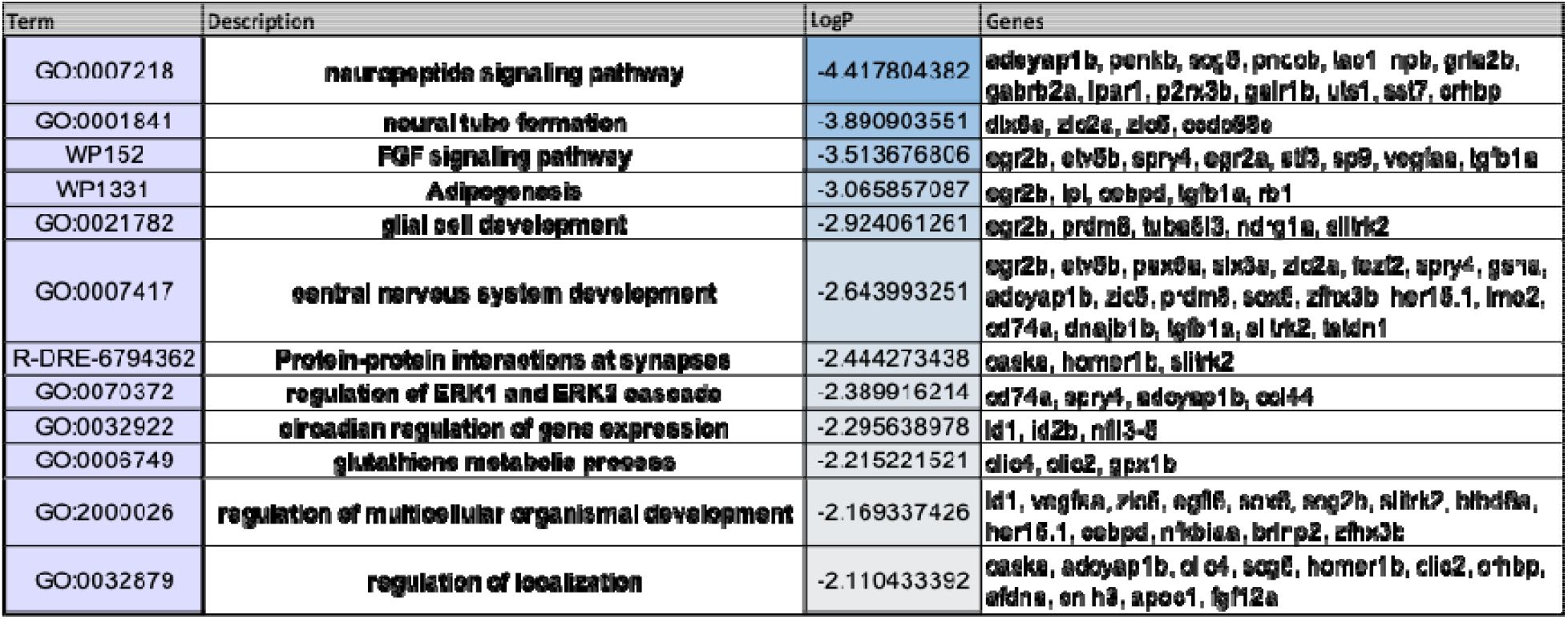
Enrichment table of the top gene ontology terms and the genes differentially expressed in neurons of the OB at 1 d after I.N. IHNV.

| Term | Description | LogP | Genes |
|---|---|---|---|
| GO:0007218 | **neuropeptide signaling pathway** | -4.417804382 | **adcyap1b, penkb, scg5, pnocb, tac1 npb, gria2b, gabrb2a, lpar1, p2rx3b, galr1b, uts1, sst7, crhbp** |
| GO:0001841 | **neural tube formation** | -3.890903551 | **dlx5a, zic2a, zic5, ccdc88c** |
| WP152 | **FGF signaling pathway** | -3.513676806 | **egr2b, etv5b, spry4, egr2a, atf3, sp9, vegfaa, tgfb1a** |
| WP1331 | **Adipogenesis** | -3.065857087 | **egr2b, lpl, cebpd, tgfb1a, rb1** |
| GO:0021782 | **glial cell development** | -2.924061261 | **egr2b, prdm8, tuba8l3, ndrg1a, slitrk2** |
| GO:0007417 | **central nervous system development** | -2.643993251 | **egr2b, etv5b, pax6a, six3a, zic2a, fezf2, spry4, [illegible], adcyap1b, zic5, prdm8, sox5, zfhx3b her15.1, lmo2, cd74a, dnajb1b, tgfb1a, slitrk2, [illegible]** |
| R-DRE-6794362 | **Protein-protein interactions at synapses** | -2.444273438 | **caska, homer1b, slitrk2** |
| GO:0070372 | **regulation of ERK1 and ERK2 cascade** | -2.389916214 | **cd74a, spry4, adcyap1b, ccl44** |
| GO:0032922 | **circadian regulation of gene expression** | -2.295638978 | **id1, id2b, nfil3-5** |
| GO:0006749 | **glutathione metabolic process** | -2.215221521 | **clic4, clic2, gpx1b** |
| GO:2000026 | **regulation of multicellular organismal development** | -2.169337426 | **id1, vegfaa, zic5, egfl6, sox8, scg2b, slitrk2, hbegfa, her15.1, cebpd, nfkbiaa, brinp2, zfhx3b** |
| GO:0032879 | **regulation of localization** | -2.110433392 | **caska, adcyap1b, clic4, scg5, homer1b, clic2, crhbp, afdna, cnih3, apoc1, fgf12a** |

